# Molecular attributes of intrinsically disordered regions in secretomes influence fungal pathogenesis

**DOI:** 10.1101/2025.10.01.679754

**Authors:** Ankita Mohapatra, Rajashekar Varma Kadumuri, Ashwin Sharma, Anita P Saju, Harikrishnan Ramadasan, Anushka Agrawal, Vaidehi Sharma, Sreenivas Chavali

## Abstract

Secretory proteins are crucial for establishing fungal infection through biomolecular interactions with the host. Accumulating evidence suggest that intrinsically disordered regions (IDRs) in proteins facilitate molecular interactions. How do IDRs of secretory proteins influence fungal pathogenesis? By analyzing 195,359 secretory proteins of 73 fungal species, we find that IDRs in plant pathogen extracellular non-effectors are enriched for weak polyampholytes and polyelectrolytes. These could adopt beads-on-string conformation effectively promoting assembly of diverse enzymes aiding swift degradation of host cell wall. Both animal pathogen secretory proteins and intracellular plant pathogen effectors show enrichment for strong polyampholytes, potentially aiding phase-separation and organizing intracellular biological matter to hijack or suppress host machinery. Importantly, IDRs of plant pathogen effectors which mimic host IDRs could have emerged through convergent evolution, while those of non-effectors might have evolved *de novo*. Thus, specific molecular attributes of fungal secretome IDRs can influence initiation, establishment and long-term persistence of infection.

## Introduction

Fungal kingdom encompasses numerous pathogens which can infect both plants and animals. Among all the plant infectious diseases, about 83% are caused by fungi [1]. Fungal pathogens have catastrophic effects on crop yield, quality and agro-economy. For instance, the rust diseases of wheat and other grain crops caused by fungal pathogens such as *Blumeria graminis, Podosphaera sps.* etc., damage about 30-40% of the crop yield [1]. Fungi also infect plethora of animals. Nearly 13 million infections and about 1.5 million human deaths annually are caused by fungal pathogens [2]. Interestingly, most of the animal fungi are opportunistic pathogens and primarily infect immunocompromised individuals, while very few are obligatory pathogens [3, 4]. Establishment of successful infection in plant and animal hosts involves multiple steps **[Supplementary Fig. 1A]**. In this context, fungal secretory proteins play essential roles by facilitating fungal colonization and morphological transitions, manipulating host cell physiology and defense [5–10].

Pathogens and hosts are constantly engaged in an evolutionary race to outsmart each other. In this context, intrinsically disordered regions/proteins (IDRs/IDPs), can serve as substrates for accelerated evolution, owing to a lack of structural constraints [11–15]. From a functional standpoint, two major features of IDRs/IDPs, viz., binding promiscuity and plasticity, facilitate their ability to bring about multitudinous molecular interactions [15–17]. Individual molecule-based studies in bacteria and oomycetes highlight that IDRs in pathogen secretory proteins could play a vital role in infection [18–21]. However, a systems-level understanding of how the functionally versatile IDRs/IDPs of fungal secretomes influence pathogenesis is still lacking. In this study, we investigated 73 fungal species and response of 16 hosts (both plants and animals) upon fungal infection to elucidate the nature and functional roles of IDRs in plant and animal fungal pathogen secretomes.

## Results

We have undertaken investigations to delineate the impact and nature of IDRs in the pathogens as well as hosts that are engaged during infection. Through an extensive literature search, we identified 124 fungal species, and selected those with well-defined pathogenicity status, and excluded fungi, whose pathogenicity status was ambiguous and those with too large (>50000 proteins) or small or partial proteomes (≤1500 annotated proteins) **[Supplementary Table 1]**. This led to the final dataset of 73 fungal species (with divergence time of ∼764 mya; **Supplementary Fig. 2; Supplementary Table 2**), of which 21 were designated as non-pathogens, 30 as plant pathogens, and 22 as animal pathogens. This classification of fungal species showed lineage specificity, with different classes of fungi overrepresented in distinct taxonomical classes **[Fig. 1A; Supplementary Table 2]**.

**Fig. 1.**
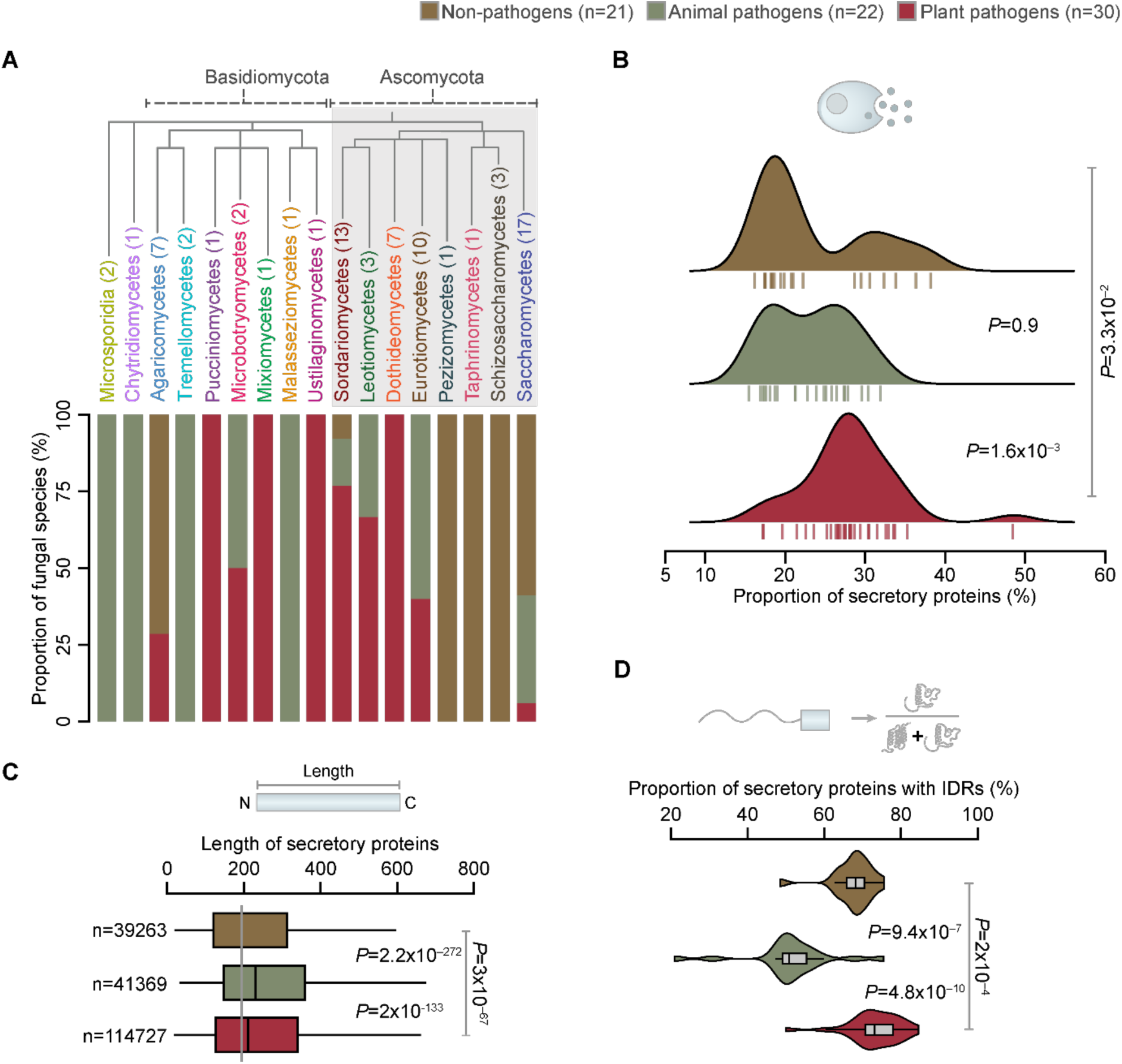
Plant pathogens contain more intrinsically disordered secretory proteins. **(A)** Distribution of fungal species based on pathogenicity (non-pathogens, animal pathogens and plant pathogens) across different fungal taxa. The numbers in the parentheses represent the total number of species of a particular fungal taxon considered in this study. Plant pathogens preponderantly belong to Sordariomycetes and Dothideomycetes, while majority of animal pathogens belong to Eurotiomycetes and non-pathogens to Saccharomycetes and Schizosaccharomycetes. These fungal taxa show distinct attributes that could contribute to this variability in pathogenesis **[Supplementary Fig. 1B]**. Interestingly, angiosperms are the most common plant hosts and mammals are the prevalent animal hosts **[Supplementary Fig. 1C-1D]**. The observed prevalence of certain taxa of the hosts might be a result of reporting bias in the literature or the pathogens with proteomes available for undertaking this study **[Supplementary Table 2]**. Nevertheless, evolutionary analysis of the plant host preferences of the fungal pathogens highlight that they could have radiated long after the emergence of angiosperms than coemerged with the origins of angiosperms [99]. **(B)** Ridgeline plot showing the distribution of the fraction of secretory proteins among non-pathogens, animal and plant pathogens. The area of the curves represents the density of the distribution for each fungal class. The bars at the bottom of the density distribution represent the data points. **(C)** Boxplot showing differences in the distribution of the length of secretory proteins, non-pathogens, animal and plant pathogens. Median of the distribution is represented by the solid line inside each box while the box limits represent the first and third quartile of the distribution. The whiskers on each end of the box show data points up to 1.5 times the interquartile range. Outliers have not been shown to facilitate better visualization. n indicates the total number of proteins in each fungal class. **(D)** Violin plot shows the fraction of proteins having intrinsically disordered regions or disordered secretory proteins among non-pathogens, animal and plant pathogens. Statistical significance was estimated using the Wilcoxon rank sum test. The number (n) provided in the parentheses in the color legend indicates the species count in each fungal class.

### Disordered secretory proteins are enriched in various biological processes that facilitate pathogenesis

We defined the fungal secretomes by predicting both classical and non-classical secretory proteins and subsequently predicted the disordered regions in these secretomes (See **Methods**). To delineate the role(s) of IDRs of fungal secretomes in pathogenesis, we drew comparisons between non-pathogens, animal pathogens and plant pathogens. Plant pathogen proteomes contain significantly higher proportion of secretory proteins while animal pathogens encoded for longer secretory proteins **[Fig. 1B-1C; Supplementary Table 2]**. Since disordered regions of length 5-10 residues are known to have functional roles, we chose a cut-off of ≥ 10 residues to define IDRs [15, 22–25]. Plant pathogen secretomes have higher fraction of proteins with IDRs compared to animal pathogens and non-pathogens **[Fig. 1D; Supplementary Table 3]**. These findings imply that enhanced intrinsic protein disorder could be a key attribute of plant pathogen secretory proteins to invade plant hosts.

Gene ontology enrichment analysis revealed that secretory proteins of all the fungal classes are involved in fundamental processes such as carbohydrate metabolism, cell wall organization and translation. Additionally, secretory proteins of plant pathogens are involved in (i) plant cell wall degradation involving metabolism of arabinan, arabinose, xylan, cellulose, glucan, and pectin and (ii) pathogenesis-related processes such as biosynthesis of toxins and metabolism of cellular aromatic compounds, organic substances and superoxides **[Fig. 2A; Supplementary Fig. 3A; Supplementary Tables 3-4]**. Contrarily, secretory proteins of animal pathogens are associated with (i) biofilm formation, which protects the fungi from host immune response and also provides drug resistance, (ii) allergic response where secretory proteins behave as Pathogen-Associated Molecular Patterns (PAMPs), leading to IgE antibody binding and triggering immune response [26, 27] and (iii) enzymes that process plant cell wall constituents such as xylan, pectin and glucan, similar to plant pathogens. Strikingly, a large proportion of secretory proteins with IDRs are primarily involved in processes that facilitate host-type specific pathogenesis **[Fig. 2A; Supplementary Fig. 3A; Supplementary Tables 3-4]**.

**Fig. 2.**
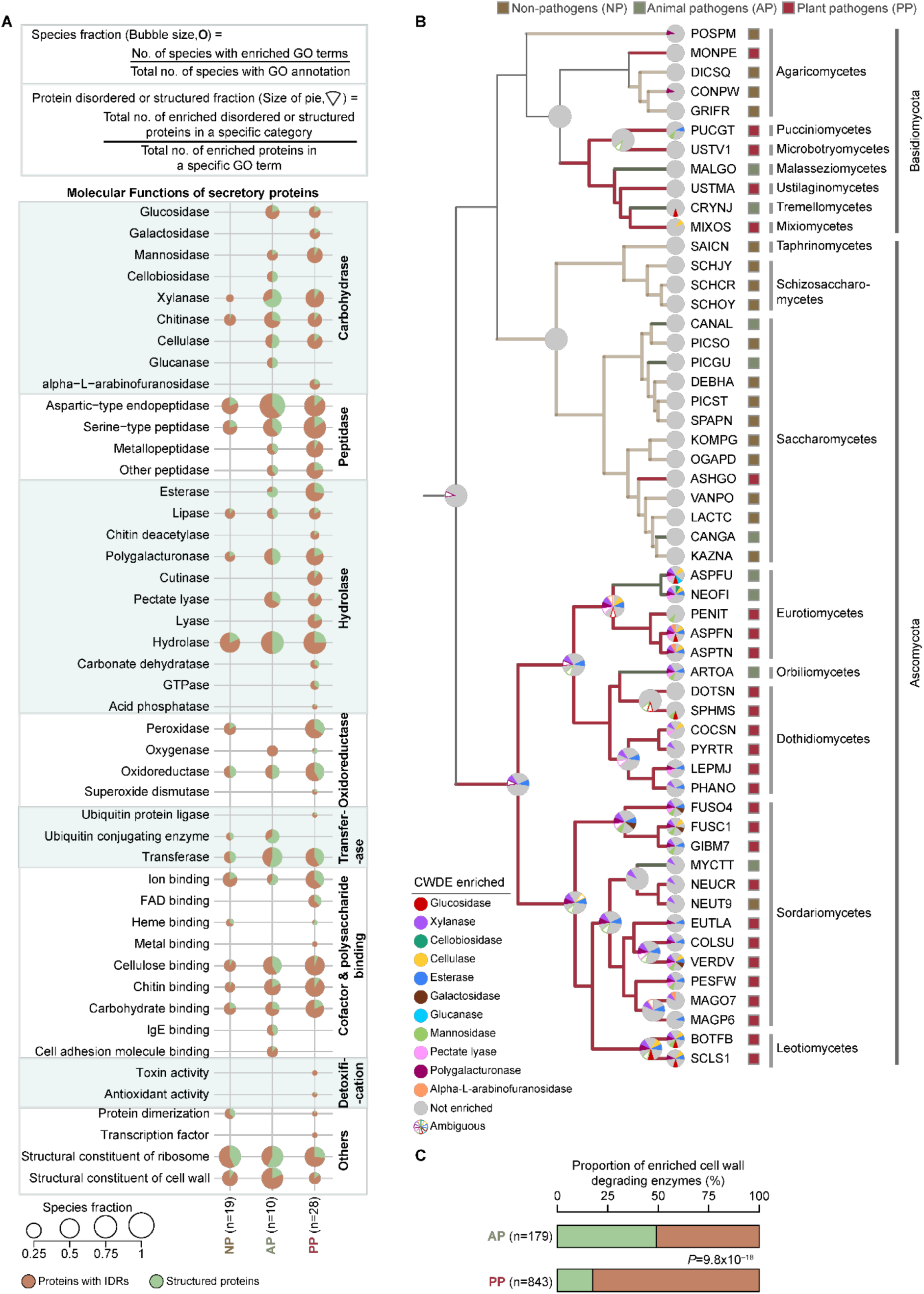
Plant pathogens contain higher fraction of intrinsically disordered cell wall degrading enzymes. **(A)** Bubble plot depicts the enriched Gene Ontology Molecular Functions (FDR < 0.05), manually grouped into different broad terms. The bubble size represents the number of species showing enrichment for a specific function. The pie chart depicts the proportion of structured and disordered secretory proteins. Proteins that contain IDR(s) were classified as proteins with IDRs while those that lack them were classified as structured proteins. Species fraction and protein disordered/structured fraction for each enriched GO term were estimated as provided in the top panel. **(B)** Phylogenetic tree depicting ancestral state reconstruction of 54 fungal species proteomes for which GO annotation was available in the DAVID server [88]. Phylogenetic tree was constructed using ProtSpaM [89]. The colour of the branches indicate the ancestral pathogenicity state reconstructed using the PastML tool [90]. The color of the pie represents the enrichment of specific cell wall degrading enzymes (CWDE) for each species and their ancestors. Ambiguous nodes are shown as white pies with the colored border, whereas nodes that are not enriched for a particular CWDE are shown using grey color. The species names have been abbreviated and the taxonomic and phylogenetic details of the species have been provided in **Supplementary Table 2**. The boxes adjacent to the species names depict the pathogenicity status (non-pathogen, animal or plant pathogen). **(C)** Bar plot showing the fraction of CWDEs which are structured and those that contain IDRs among animal and plant pathogens. Statistical significance was estimated using Fisher’s exact test.

Why are CWDEs enriched in animal pathogens? To address this, we performed ancestral state reconstruction analyses of 54 fungi and identified two major lineages– a non-pathogen and a mostly plant pathogen lineage. Two-thirds of the animal pathogens correspond to the plant pathogen lineage. Enrichment of CWDEs was preponderantly observed in both plant pathogens and animal pathogens of the plant pathogen lineage. These results highlight that CWDEs could have evolved from a common ancestor during fungal evolution **[Fig. 2B]**. In the absence of appropriate hosts (often immunocompromised individuals) most of the animal pathogens behave as saprotrophs, which survive largely on plant debris [28–31]. This explains the enrichment of plant cell wall degrading enzymes (CWDEs) in their secretomes. On the contrary, plant pathogens that have emerged in non-pathogen lineage lack CWDEs and penetration plugs. Such pathogens infect plants (i) with the help of insect vector (*Ashbya gossypii* [32]), (ii) by penetrating through large stomatal pores (*Dothistroma septosporum* infection of conifers [33]), or (iii) by replacement of pollen grains with fungal spores, which are transmitted by pollinating insects (anther smut, *Microbotyum lychnidis-dioicae* [34]). Strikingly, plant pathogen CWDEs contain significantly more proteins with IDRs than those of animal pathogens (83% versus 51%) **[Fig. 2C; Supplementary Tables 3-4]**. Among the shared enriched biological processes, plant pathogens tend to show a higher proportion of proteins with IDRs **[Supplementary Fig. 3B**-3D**; Supplementary Tables 3-4]**. While CWDEs are known to have expanded in the Dikarya lineage (both ascomycota and basidiomycota species) [35], our findings highlight higher acquisition of IDRs within CWDEs of plant pathogen secretomes, implying that they may play an important role in their pathogenesis.

### Plant and animal pathogen secretomes show distinct nature of disordered regions which might influence their infectability

The location and composition/conformation of disordered segments (referred henceforth as state IDRs) affect functionality [15, 36]. Do the IDRs in the secretomes of fungal pathogens exhibit properties that may influence pathogenicity? To examine location bias of IDRs among the secretomes of the three classes of fungi, we binned the protein into N-terminal, Internal and C-terminal regions, using tertile cut-offs. Based on their location, IDRs were assigned as N-terminal, Internal, C-terminal, N–Internal, Internal–C and N–C **[Fig. 3A; Supplementary Table 5]**. While secretomes of plant pathogens are enriched for N-terminal and N–C IDRs, those of animal pathogens are enriched for C-terminal IDRs. Contrarily, non-pathogenic secretomes are enriched for Internal IDRs **[Fig. 3A; Supplementary Table 5]**.

**Fig. 3.**
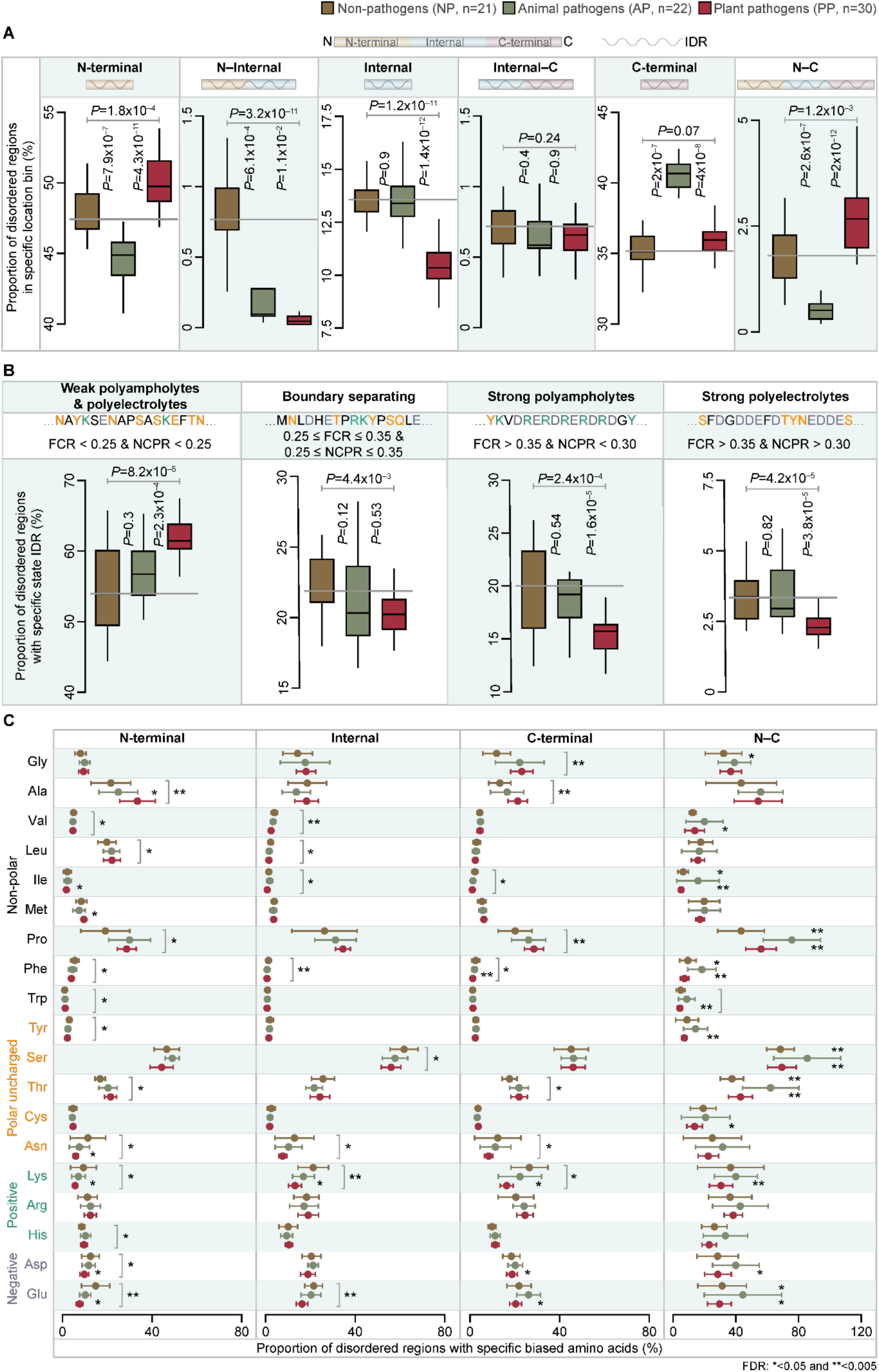
Different classes of fungi show distinct bias in the location and composition of secretome IDRs. **(A)** Boxplot showing the distribution of disordered regions across different location bins in secretory proteins of different classes of fungi. Based on length tertiles, each fungal secretory protein was classified into three regions: N-terminal, Internal and C-terminal. We binned IDRs based on their location, across these three regions into six categories as (i) N-terminal, Internal, C terminal, if > 50% of their length aligned to one of the three regions, (ii) N–Internal, Internal–C if the IDR was distributed equally across the two regions and (iii) N– C, if the entire protein was predicted to be an IDP. **(B)** Boxplot showing the distribution of different state IDRs of secretory proteins across the three fungal classes. The top part of the panel highlights the classification of IDRs into different states based on the bias for charged residues estimated by (i) Fraction of Charged Residues (FCR) which represents the sum of the fraction of positively charged residues (f^+^) and negatively charged residues (f^−^) and (ii) Net Charge Per Residue (NCPR) i.e., difference in the f^+^ and f^−^, IDRs. These include (i) Weak polyampholytes and polyelectrolytes (FCR and NCPR < 0.25), (ii) Boundary separating (FCR and NCPR ≥ 0.25 and ≤ 0.35), (iii) Strong polyampholytes (FCR > 0.35 and NCPR < 0.30) and (iv) strong polyelectrolytes (FCR > 0.35 and NCPR > 0.30) [37]. Statistical significance was obtained by Wilcoxon rank sum test. The number of datapoints in each box plot in panels (**A**) and (**B**) correspond to the number of species in each class. **(C)** Distribution of biased amino acids, estimated using FLPS [93], in the IDRs across different protein location bins. Circles represent the median values and bars represent Median Absolute Deviation (MAD). The colors indicate the class of fungi. Statistical significance was estimated using Wilcoxon rank sum test, and asterisks near the bars indicate the False Discovery Rates (FDR) computed using Benjamini-Hochberg method of correction for multiple testing.

Based on the bias for charged residues estimated by Fraction of Charged Residues (FCR) and Net Charge Per Residue (NCPR), IDRs can be classified as weak polyampholytes and polyelectrolytes, boundary separating, strong polyampholytes and strong polyelectrolytes [37]. Secretomes showed distinct enrichment of (i) weak polyampholytes and polyelectrolytes in plant pathogens, (ii) strong polyampholytes and strong polyelectrolytes in animal pathogens and (iii) all state IDRs barring weak polyampholytes and polyelectrolytes in non-pathogens **[Fig. 3B; Supplementary Table 6]**. These findings are not confounded by the threshold of IDR length (10 amino acids) considered here **[Supplementary Fig. 4A**-4B**; Supplementary Tables 5-6]**. Furthermore, IDRs in secretomes of different classes of fungi show distinct amino acid compositional bias **[Fig. 3C; Supplementary Table 7]**– (i) non-polar (Ala, Leu, Met) and polar uncharged (Thr) residues in N-terminal IDRs in plant pathogens facilitating hydrophobic interactions, (ii) polar residues (Ser, Lys, Asp, Glu) and Pro in C-terminal and N–C IDRs of animal pathogens aiding charge-based interactions and (iii) polar uncharged (Ser and Asn) and charged residues (Lys and Glu) in the internal IDRs of non-pathogens promoting coiled-coil and charge-based interactions. Moreover, we could recapitulate the trends associated with enrichment of IDRs and their class-specific attributes in fungal secretomes, across different orthologous groups [38] of secretory proteins **[Supplementary Fig. 5; Supplementary Table 8]**. Thus, even within homologous proteins, IDR-specific attributes might have been acquired and retained due to their influence on pathogenesis.

Plant pathogen secretory proteins with weak polyampholytes and polyelectrolytes are enriched for metabolism of various type of carbohydrates and their respective degradative enzymes like glucosidase, galactosidase, mannosidase and xylanase **[Supplementary Fig. 6; Supplementary Table 4]**. Weak polyampholyte and polyelectrolytes aid formation of beads on string-like structures providing multivalency to a protein, which promotes simultaneous multi-partner binding [39, 40]. We posit that such beads across different plant pathogen secretory proteins, may facilitate (i) effective adherence of the proteins to cell wall and/or (ii) rapid organization of multiple units of the same protein or different carbohydrate degrading enzymes, simultaneously in the extra-cellular space to aid swift degradation of the cell wall and quick penetration of the host cell.

Contrarily, animal pathogens do not require cell wall degradation for their infection, suggesting that most secretory proteins might enter the host cell and modulate host cellular processes. Electrostatic interactions driven by strong polyampholytes and strong polyelectrolytes bring about moderate specificity and moderate affinity-based interactions, that help engage different host proteins [41, 42]. In addition, bias for charged residues (Lys, Glu and Asp) in the C-terminus of animal pathogens might contribute to the enhanced stability of the pathogenic proteins inside the animal host [36, 43, 44]. Collectively our findings indicate a distinct location, nature and amino acid bias in IDRs of the secretomes of plant and animal pathogens, with implications to their infectability.

### Plant pathogen effectors and non-effectors show distinct IDR attributes that facilitate intracellular and extracellular organization of biological matter

During plant infections, pathogen effector proteins tend to enter the host and manipulate host immunity, while certain secretory proteins remain preponderantly extra-cellular and aid cell wall degradation (referred henceforth as non-effectors). How do the IDR attributes of secretory proteins that remain extra-cellular differ from those that enter the host cells? We classified plant pathogen secretory proteins into those that are predicted to be effectors and others as non-effectors and compared their IDRs. Effectors are significantly shorter and contain fewer proteins with IDRs than non-effectors **[Fig. 4A-4B; Supplementary Table 3]**. Interestingly,

i. non-effectors are mostly localized to the outer surface of the cell and involved in metabolism of different substrates related to cell wall degradation and nutrient uptake, whereas (ii) effectors are localized to multiple intra-cellular compartments and enriched for intracellular processes such as transcription, translation and cell cycle **[Fig. 4C; Supplementary Fig. 7; Supplementary Table 4]**. Our observations corroborate with previous studies [45–47]. While these enrichments are pathogen-based, they reinforce the reliability of the classification of effectors and non-effectors as these secretory proteins might have similar localizations in the hosts, as well.

**Fig. 4.**
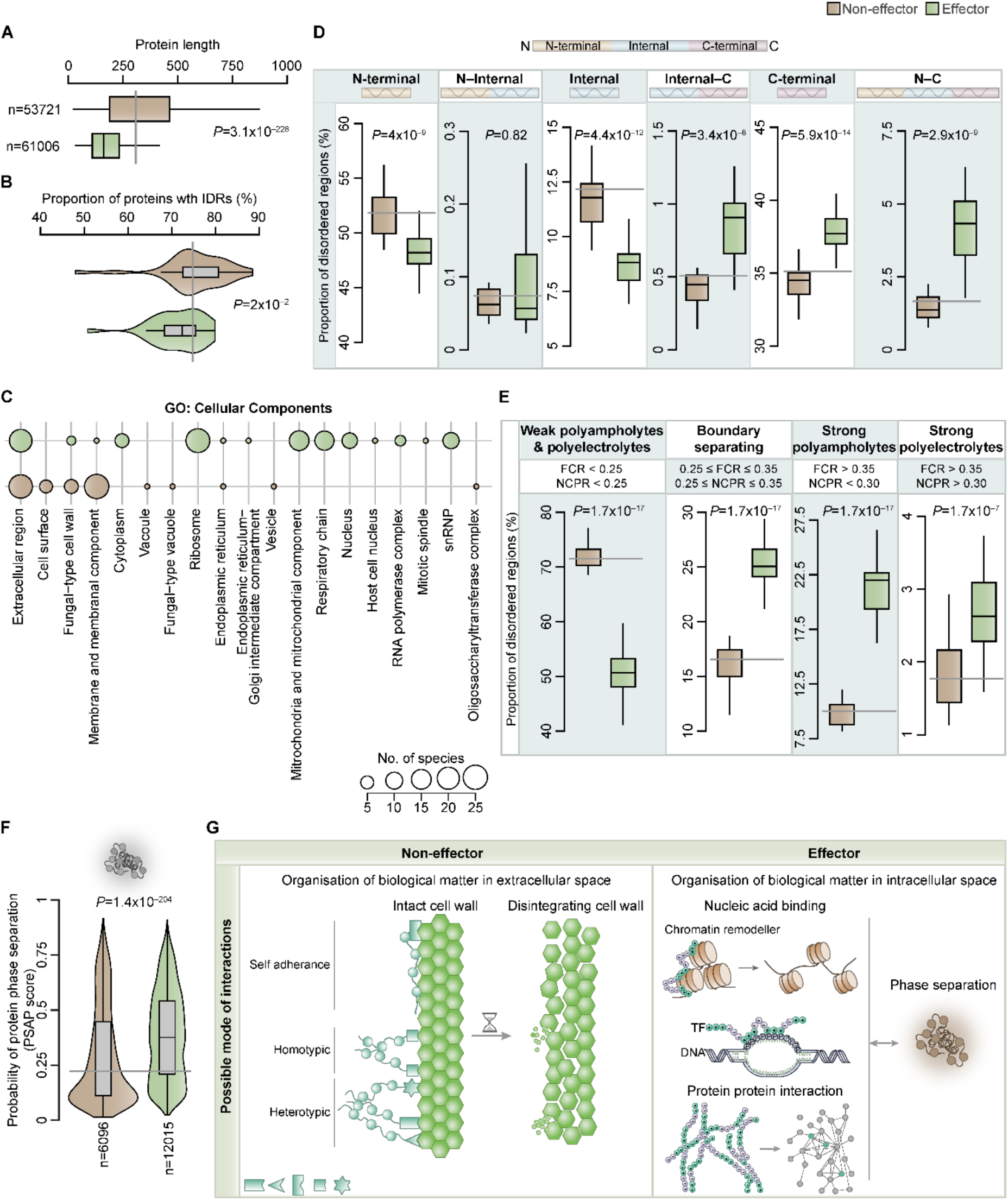
Effectors and non-effectors of plant pathogens show distinct IDR attributes that could aid organization of biological matter for specific functionalities in distinct locations. **(A)** Box plot showing the distribution of protein length of effectors and non-effectors of plant pathogens. **(B)** Violin plot shows the fraction of proteins with IDRs among effectors and non-effectors. **(C)** Bubble plot depicts the enriched GO-Cellular Components; manually classified into broad categories. The bubble size denotes the number of species representing enriched GO terms in each category. Boxplot showing the distribution of IDRs in effectors and non-effectors **(D)** across different regions in proteins classified based on length tertiles and **(E)** the different states of IDRs. **(F)** Violin plot showing the probability of phase-separated structure formation, represented by the PSAP (Probability of protein phase separation) scores [56], for proteins containing strong polyampholyte IDRs, among effectors and non-effectors. **(G)** Schematic representation of how distinct IDR attributes in non-effectors and effectors facilitate organization of biological matter in extra-cellular and intra-cellular space respectively to affect different processes during infection.

While non-effectors are enriched for N-terminal and internal IDRs, effectors showed bias for C-terminal and N–C IDRs. From the state IDR stand point, (i) non-effectors are enriched for weak polyampholytes and polyelectrolytes with bias for non-polar and polar uncharged residues (Ser, Thr and Cys) whereas (ii) effectors showed bias for strong polyampholytes and polyelectrolytes and boundary separating IDRs with bias for polar uncharged (Asn and Gln) and all charged residues **[Fig. 4D-4E; Supplementary Fig. 8; Supplementary Tables 5-7]**. Our observations pertaining to non-effectors corroborate with the previous finding that carbohydrate-binding proteins are enriched for low-complexity regions rich in serine and threonine [48]. Strikingly, weak polyampholytes and polyelectrolytes containing non-effectors predominantly contain domains associated with the metabolism of carbohydrates, proteins, and lipids **[Supplementary Fig. 9A; Supplementary Table 9]**. Thus, the presence of N-terminal weak polyampholytes and polyelectrolytes in non-effectors forming potential beads on string-like structures might facilitate organization of biological matter in the extra-cellular space for swift degradation of plant cell wall and nutrient uptake by the pathogen.

Conversely, C-terminal strong polyampholytes and strong polyelectrolytes in effectors might aid the formation of phase-separated condensates [49–52], through electrostatic interactions [53–55], for altering or hijacking host cellular processes to aid pathogenesis. Remarkably, among strong polyampholytes-containing proteins, effectors show higher probability of phase separation [56] than non-effectors **[Fig. 4F; Supplementary Table 9]**. Examination of charge patterning (estimated as kappa; κ) revealed a greater tendency for the strong polyampholytes in effectors to form higher number of interchain interactions, which facilitate the formation of phase-separated condensates **[Supplementary Fig. 9B; Supplementary Table 9]** [54, 57]. Besides charge patterning, post-translational modifications (PTMs) are important contributors of phase separation [58]. Effectors tend to contain higher PTM density than their non-effectors counterparts **[Supplementary Fig. 9C; Supplementary Table 9]**. Collectively, effectors have various attributes that promote organization of intracellular biological matter.

To ascertain our observations, we collated experimentally determined pathogenesis-related proteins from PHI-base [59] and classified them into effectors and non-effectors, as done for the secretomes. We observed that the effectors (i) are shorter, (ii) contain fewer proteins with IDRs, (iii) have more boundary separating and strong polyampholyte IDRs and (iv) have higher C-terminal disorder. Akin to our aforementioned observations, effectors among the experimentally determined proteins were predominantly involved in transcription and post-transcriptional modification, translation and nucleic acid binding **[Supplementary Fig. 10; Supplementary Table 10]**. Taken together, IDRs have emerged to serve distinct functionalities in different spatial localizations in terms of organizing biological matter (i) extracellularly for non-effectors to rapidly degrade host cell wall and (ii) intracellularly for effectors to regulate various host cellular processes, thereby contributing to fungal infections **[Fig. 4G]**.

### IDRs of fungal pathogen secretomes mimic the host infection-response proteins

Since pathogens are known to mimic host proteins to suppress or hijack host machinery or processes [60, 61], we investigated the extent of overlap between IDRs of fungal pathogen secretomes and the host infection-response proteins to decipher plausible evolutionary trajectories. To delineate the influence of IDRs on host infection-response, we assembled and investigated 24 publicly available host transcriptomic datasets corresponding to 14 different plants infected by 13 plant pathogens and 2 animals infected by 5 animal pathogens **[Fig. 5A; Supplementary Table 11]**. For this, we examined the IDR attributes of proteins resulting from translation of differentially expressed host transcripts upon infection (referred henceforth Differentially Expressed Proteins; DEP). We find a higher proportion of proteins with IDRs among the DEPs of both plant and animal hosts **[Fig. 5B; Supplementary Table 12]**. Next, we estimated the extent of correlation between the fraction of over-represented state IDRs in the fungal pathogens with those in DEPs of the corresponding hosts. We could not detect any correlation between the strong polyampholytes and strong polyelectrolytes of animal DEPs with those of animal pathogen secretory proteins **[Fig. 5C; Supplementary Tables 13-14]**. This could be probably due to fewer available animal host transcriptomic datasets or a lesser likelihood of substantially different IDR attributes of animal pathogens compared to animals. Conversely, plant pathogen effectors showed positive correlation for strong polyampholytes with plant DEPs **[Fig. 5C; Supplementary Tables 13-14]**. The usage of host-like strong polyampholyte containing IDRs, especially for gene-expression regulation, implies that plant pathogen effectors might have acquired molecular mimicry either through horizontal gene transfer or convergent evolution. Sequence identity-based comparison of fungal strong polyampholytes with that of host DEPs shows that pathogen strong polyampholytes might have predominantly emerged through convergent evolution **[Supplementary Fig. 11]**. However, we did not find any correlation between the weak polyampholytes and polyelectrolytes containing IDRs of the pathogen non-effectors with those of hosts **[Fig. 5C; Supplementary Tables 13-14]**. This suggests that the preponderance of weak polyampholytes and polyelectrolytes containing IDRs in the non-effectors, facilitating assembly of CWDEs for rapid degradation of host cell wall, might have emerged as a novel IDR attribute specific to plant pathogens. Taken together, IDRs in effectors that mimic those of the host regulators might have emerged predominantly through convergent evolution, whereas those of non-effectors might have evolved *de novo* aiding rapid organization of CWDEs leading to rapid degradation of host cell wall **[Fig. 5D]**.

**Fig. 5.**
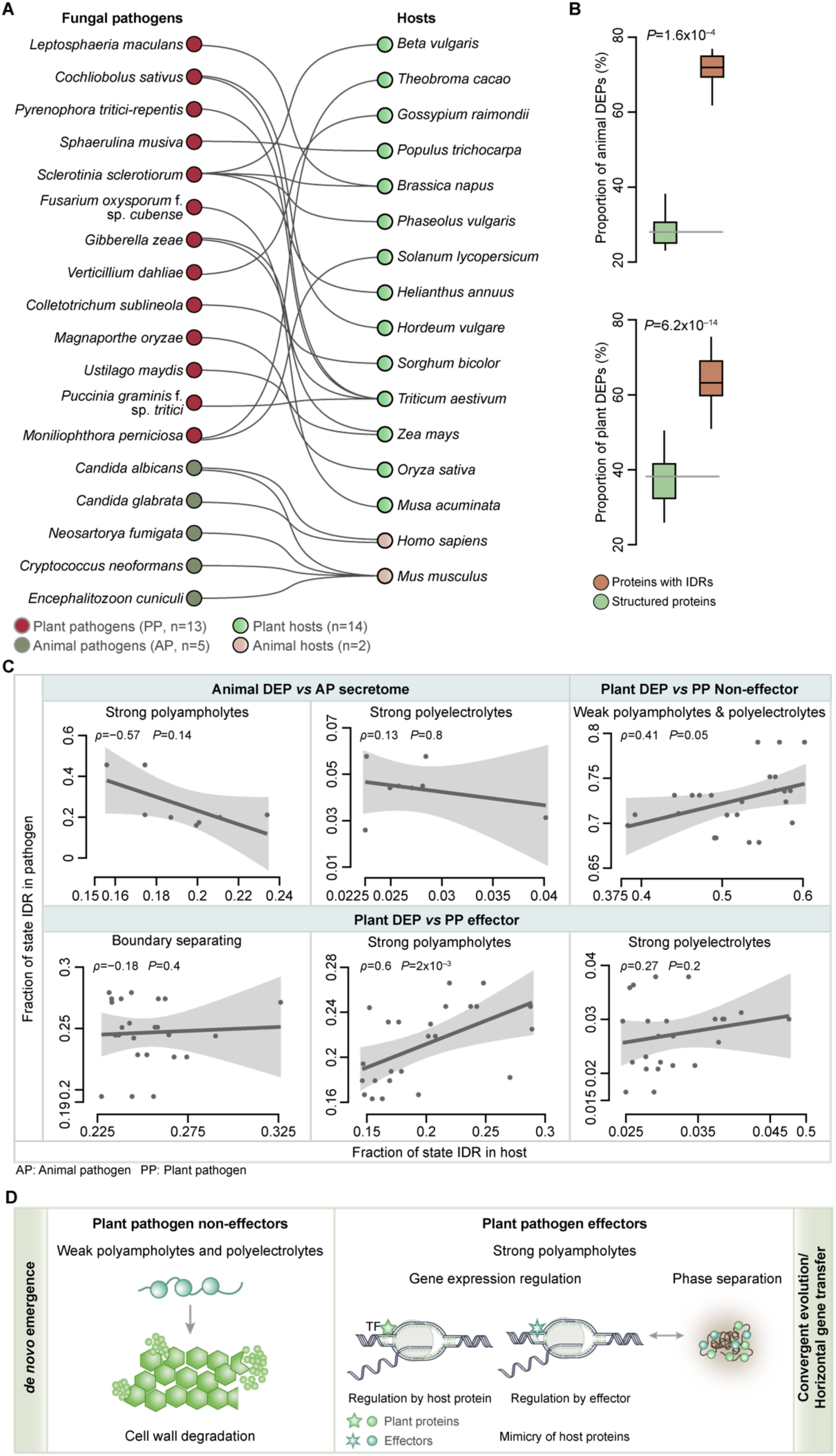
Possible modes of emergence of IDRs of fungal pathogen secretomes. **(A)** Bipartite network of the transcriptomic datasets of hosts (shown on the right) infected by fungal pathogens (shown on the left) analyzed in this study. The color of the nodes corresponds to the different types of hosts and pathogens. **(B)** Box plot of distribution of proportion of proteins with IDRs among differentially expressed proteins (DEPs) in both animal and plant hosts upon fungal infection. **(C)** Line plot depicting the correlation between the fraction of overrepresented state IDRs in the animal and plant pathogen (both effectors and non-effectors) secretomes with the fraction of corresponding state IDRs in the respective host DEPs. Significance of correlation was estimated using Spearman rank correlation. The grey band around the correlation trend line represents confidence interval (95% CI) while ‘*ρ*’ and ‘*P*’ denote the Spearman’s rank correlation coefficient and the statistical significance estimate, respectively. **(D)** Schema representing possible evolutionary trajectories of the weak polyampholytes and polyelectrolytes of plant pathogen non-effectors and strong polyampholytes of plant pathogen effectors with respect to the host IDRs.

## Discussion

The current understanding of the role of IDRs in infections is largely based on molecular studies of individual proteins or IDRs of bacteria, viruses and oomycetes [19, 20, 62, 63]. In fungi, molecular studies thus far have deciphered the role of IDRs in development [64–66] and stress response [67, 68]. However, how IDRs in fungi influence pathogenicity remains largely unknown. From a functional stand point we show that (i) plant pathogen non-effectors are enriched for weak polyampholytes and polyelectrolytes IDRs at the N-terminus, which could aid rapid assembly of diverse CWDEs in the extra-cellular space, (ii) plant pathogen effectors and animal pathogen secretory proteins are enriched for strong polyampholytes at the C-terminus aiding **[Fig. 6]** organization of intracellular biological matter, thereby promoting hijacking or suppressing host gene expression. From an evolutionary standpoint, our observations revealed a possible *de novo* emergence of weak polyampholytes and polyelectrolytes of plant pathogen non-effectors and a predominantly convergent mode of evolution of the host-like strong polyampholytes in plant effectors **[Fig. 5D]**.

**Fig. 6.**
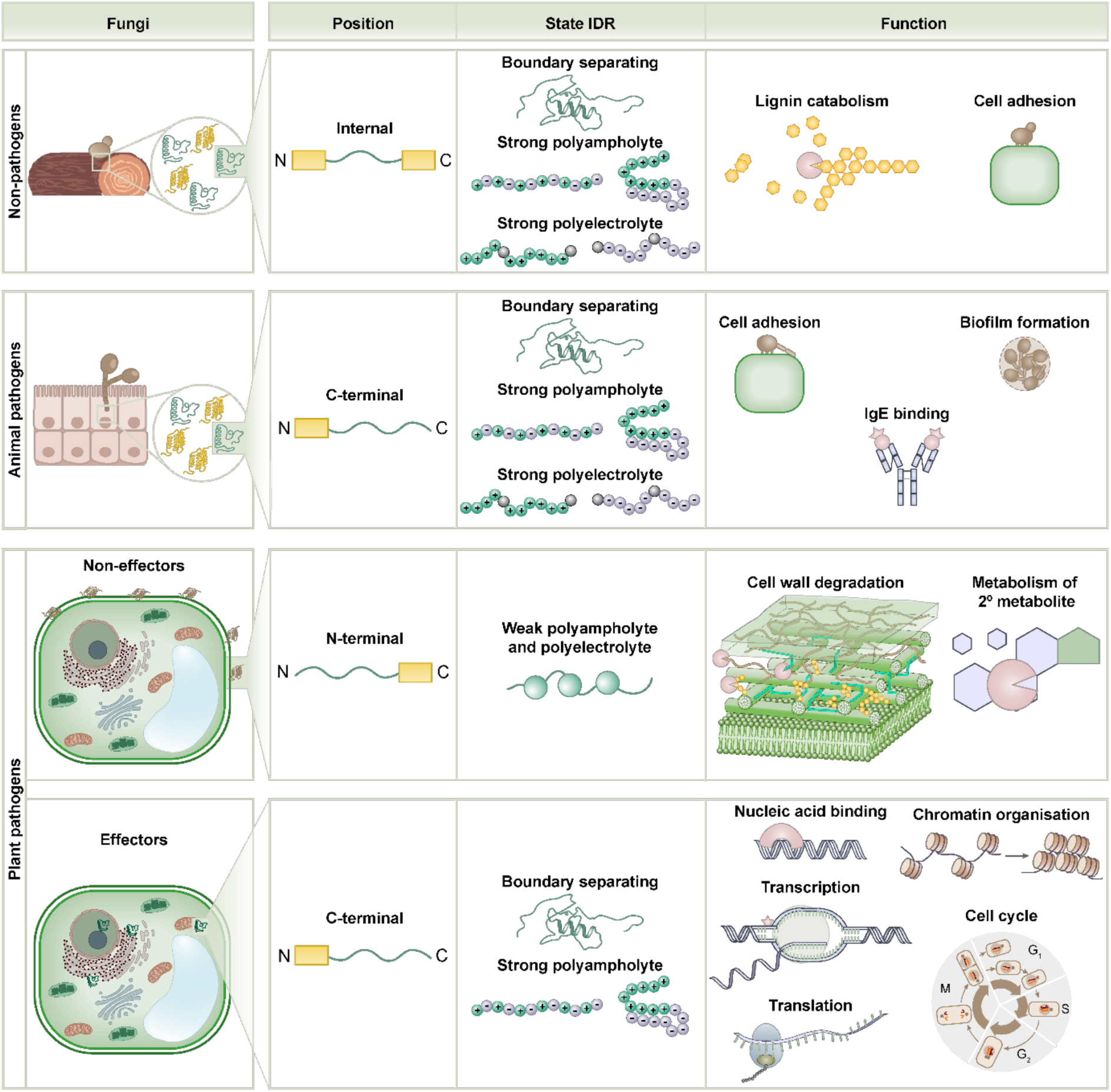
Prevalence and consequences of distinct attributes of secretome IDRs in fungal pathogens. Non-pathogen secretome IDRs are found predominantly in internal regions of proteins and show bias for strong polyampholytes and polyelectrolytes, which might influence lignin metabolism and cell adhesion (first panel). Animal pathogen secretomes show preponderance for C-terminal disorder biased for strong polyampholytes and strong electrolytes, potentially influencing processes such as biofilm formation and IgE binding (second panel). While plant pathogens non-effectors are biased for N-terminal IDRs enriched for weak polyampholytes and polyelectrolytes influencing functions such as cell wall degradation (third panel), plant pathogen effectors show preferential enrichment for strong polyampholytes and polyelectrolytes in the C-terminal regions influencing host regulation (fourth panel).

IDRs in fungal secretomes may have additional roles to those identified in this study. For instance, IDRs can facilitate (i) fungal penetration of host (e.g., LPMO of the plant pathogen *Colletotrichum orbiculare* [69], Aip5 of the animal pathogen *Candida albicans* [70]) (ii) effector translocation (e.g., Kex2 of *Parastagonospora nodorum* [71]) and (iii) subversion of the host defense either through IDR-mediated protein unfolding, (e.g., PWL2 of plant pathogen *Magnaporthe oryzae* [72]) or through disorder to order transition (e.g., ZCF3 of the animal pathogen *Cryptococcus neoformans* [73]). Additionally, IDRs in the secretory proteins can also affect (i) molecular promiscuity, implying that the same region can interact with multiple host or other pathogen proteins and (ii) half-life of the pathogen proteins, which can be enhanced by C-terminal IDR tails biased for charged residues [36, 43]. Importantly, the functionalities of IDRs in promoting infections show similarity across different kingdoms of pathogens. This is exemplified by the IDR-mediated (i) translocation of RxLR of the oomycete *Phytophthora sojae* [20] and (ii) evasion of host defense by Avr2 of *Phytophthora infestans*

*(i)* [62] and the phytobacterial type III effectors of *Xanthomonas campestris* [19]. IDR-mediated phase separation is known to contribute to bacterial pathogenicity. For instance, (i) effectors of *Pseudomonas syringae* form condensates to repress host translation, thereby promoting infection [74] and (ii) the multivalent IDR of the Type III effector XopR of *Xanthomonas campestris* aids condensate formation through interaction with actin-binding proteins, which enables the pathogen to evade plant defense [19]. We find that fungal effectors with strong polyampholyte IDRs exhibit a higher tendency for phase separation, suggesting that they could employ similar phase separation-mediated strategies to circumvent host defense.

IDRs exhibit unique features such as rapid sequence divergence both in terms of composition and length, ability to form conformational ensembles and diverse modes of action. These features could contribute to (i) swift acquisition of functional specialization or versatility of fungal secretomes and/or (ii) limit the ability of host immune receptors to evolve specificity against fungal pathogens. Additionally, fusion of IDRs with structured regions could lead to rapid expansion of functional repertoire of secretory proteins, as shown in the core Tin2 fold of *Ustilago maydis* [75]. Engagement of such dynamic IDRs in fungal secretomes not only represent an efficient strategy to constantly achieve one-upmanship over the host immune system but also provides an avenue to expand the host spectrum. Notwithstanding such accelerated evolution, observation of consistent patterns of IDRs affecting overlapping functions across pathogens might indicate positive selection of IDR attributes for distinct biological functions in a host-type specific manner. Focused molecular, biochemical and structural studies will provide insights on the precise mode(s) of secretome IDR engagement during infections. Given that the hosts also engage IDRs to defend pathogens [76–78] rigorous evolutionary studies are required to delineate how the dynamics of co-evolution of IDRs in host and pathogen shapes the evolutionary landscape of IDR-mediated host-pathogen interactions.

The findings and interpretations presented here might be influenced by large-scale datasets integrated and analyzed here, and the performance of the tools used. Although we have used state-of-the-art methods to define IDRs and secretory proteins and for classifying plant pathogen secretory proteins into effectors and non-effectors, the predictions might not precisely represent them. The binary classification of secretory proteins into effectors and non-effectors might be confounded by (i) our current understanding of effector properties that are employed by the predictors for effector prediction and (ii) the possibility that certain proteins might perform the functions of both classes of proteins. Furthermore, general trends observed in this study might not be applicable for each and every secretory protein in each individual fungal pathogen examined. This necessitates robust molecular as well as high throughput studies to elucidate the impact of various IDR attributes identified here on fungal pathogenesis. As shown using the vanillin derivatives which effectively inactivate the tomato spotted wilt virus (TSWV) by targeting key residues located in IDR of the viral nucleocapsid protein critical for phase separation [79], we envisage that the fungal secretome IDR attributes influencing key functions identified here, present unprecedented opportunities for developing new therapeutic strategies to tackle fungal infections.

## Methods

### Compilation and classification of fungal species

We collated about 124 fungal species with annotated proteomes in the Uniprot [Release 2021_03] [80] and with orthologs documented in the OMA orthology database [Release 2021] [38] [**Supplementary Table 1**]. Information regarding pathogenicity of the fungal species was obtained from USDA - U.S. National Fungus Collections Fungal Database (https://fungi.ars.usda.gov/) and through extensive literature search. The taxa information of each fungal species was obtained using the NCBI Taxonomy browser [81] and Tree of life (http://tolweb.org/tree/). Fungi with ambiguous annotation for pathogenicity status, including those infecting both plant and animal hosts or with too large (>50000 proteins) or small or partial proteomes (≤1500 proteins) were disregarded. This resulted in a total of 73 fungal species consisting of about 727,400 proteins **[Supplementary Table 2]**. Based on their pathogenicity status, these fungal species were categorized into animal pathogens (n=22) that infect animals including humans, plant pathogens (n=30), which infect plants and well-studied fungal species with no known information on infections were classified as non-pathogens (n=21) **[Supplementary Table 2]**.

### Prediction of secretory proteins and effectors

Fungal secretory proteins are categorized into classical and non-classical secretory proteins. Classical secretory proteins are characterized by the presence of an N-terminal secretory signal peptide. Contrarily non-classical secretory proteins, lack signal peptides and are also known as leaderless sequences. These are identified through various features such as amino acid composition, the number of atoms (C, H, N, and S), positively charged residues, pro-peptide cleavage sites, protein sorting, low complexity regions, and transmembrane helices [82]. We predicted classical secretory proteins using SignalP V 5.0 [83] with a default D-score cutoff of ≥ 0.5 and non-classical secretory proteins, which lack signal peptides, using SECRETOOL [82] and applied the default Neural Network score threshold of ≥ 0.8. The fungal secretomes analyzed in this study include both the predicted classical and non-classical secretory proteins, amounting to 195,359 secretory proteins across 73 fungal species **[Supplementary Table 3]**. We predicted a total of 61,006 plant pathogen effector proteins using the two machine learning models of EffectorP 3.0 [84], which were trained on apoplastic effectors and cytoplasmic effectors. Plant pathogen secretory proteins that were not identified as effectors were classified as non-effectors (n=53,721).

### Prediction of intrinsically disordered regions

Intrinsically disordered regions in the fungal secretory proteins and host DEPs were predicted using AUCPreD [85]. AUCPreD is amongst the top five accurate predictors of IDRs and has high computational speed that facilitates analyzing large-scale datasets, as studied here [86]. Proteins with IDRs of length ≥ 10 residues were classified as disordered proteins. Contiguous IDRs (minimum of 4 residues each, as identified by AUCpreD) separated by ≤4 amino acids (minimum number of residues required for making a secondary structure) were merged and treated as a single disordered tract using an in-house written Python script. A total of 132,102 proteins were predicted as proteins with IDRs and proteins that lacked the IDRs were classified as structured proteins **[Supplementary Tables 3 and 12]**.

### Protein domain prediction

A total of 48,420 secretory proteins containing N-terminal weak polyampholytes and polyelectrolytes including both effectors and non-effectors of plant pathogen were subjected to domain prediction analysis using Reverse Position-Specific BLAST (rpsBLAST) [87]. The rpsBLAST application searches a protein query against the conserved domain database (CDD), which comprises a comprehensive collection of curated protein domain profiles. By aligning each query sequence against these profiles, rpsBLAST enables the identification and annotation of conserved functional domains within the proteins **[Supplementary Table 9]**.

### Gene ontology enrichment analysis

Gene Ontology (GO) enrichment analysis of different classes of secretory proteins for biological processes, molecular functions and cellular components was done using the DAVID server, with a threshold set for FDR < 0.05 [88]. Enriched GO terms were then manually grouped into broader categories **[Supplementary Table 4]**.

### Ancestral state reconstruction

A species-level phylogenetic tree was generated for 54 fungal species, for which GO annotation was available in DAVID (including non-pathogens, animal pathogens and plant pathogens). Using the whole-proteome sequence of these species, we performed phylogeny reconstruction by employing ProtSpaM, an alignment-free method based on Filtered Spaced Word Matches (FSWM) [89]. The taxonomic classification (rank: class and phylum) was obtained from NCBI Taxonomy browser. Information regarding the pathogenicity status of fungal species and the enrichment of different types of cell wall degrading enzymes (CWDE), based on GO annotation information were superimposed on the tree. To examine the evolutionary trajectory of pathogenicity in these species, we performed ancestral character state reconstruction using PastML [90], using the rooted phylogenetic tree with the tips annotated with pathogenicity data. We employed the recommended ML method, marginal posterior probabilities approximation (MPPA) under F81 character evolution model [91] and iTOL (https://itol.embl.de/) for visualization and annotation of the results.

### Classification of disordered proteins

Disordered proteins were classified based on two attributes as follows.

*(i) Location of the IDRs in the protein:* Using an inhouse python script, we first binned IDRs into three protein location bins viz., N-terminal, internal, and C-terminal, identified based on the protein length tertiles. For IDRs that spanned across more than one bin, we assigned the IDR to the bin in which more than 50% of the IDR was located. IDRs evenly distributed across different protein location bins between two tertiles, were classified into overlapping bins such as N-terminal–Internal, Internal–C-terminal. Proteins that were completely disordered were classified in the N-terminal–C-terminal bin. Thus, IDRs were classified across six different protein location bins **[Supplementary Table 5]**.
*(ii) Physicochemical state of IDRs:* Sequence features of IDRs, including net charge per residue (NCPR = |f^+^ – f^−^|; where f^+^ and f^−^ indicate the fraction of positively and negatively charged residues respectively) and the fraction of charged residues (FCR = f^+^ + f^−^), were obtained using the Python module localCIDER [92]. Based on NCPR and FCR values, the IDRs in the fungal secretomes and host DEPs were classified into four groups, or states of IDR: (i) weak polyampholytes and polyelectrolytes, (ii) boundary separating, (iii) strong polyampholytes, and (iv) strong polyelectrolytes, in accordance with Das-Pappu diagram of states [37] **[Supplementary Tables 6 and 13]**. Charged patterning of strong polyampholytes of plant pathogen secretory proteins, reflected by the segregation of positive and negative charged amino acids in the IDRs was determined by estimating the kappa value using localCIDER [92] **[Supplementary Table 9]**.

### Identification of biased amino acid(s) in the IDRs

Regions exhibiting compositional bias within the IDRs were identified using fLPS 2.0 [93]. fLPS analyzes sequence composition by comparing observed versus expected frequencies of specific amino acids, with a binomial P value ≤ 0.01 **[Supplementary Table 7]**.

### Post-translational modifications

Experimentally identified post-translational modification (PTM) sites and types of effector and non-effector proteins were retrieved from dbPTM [94]. We estimated the PTM density of each secretory protein as the ratio of the number of PTM sites to the entire length of the protein **[Supplementary Table 9]**.

### Prediction of probability of phase separation

We used the tool Phase Separation Analysis and Prediction (PSAP) [56] to estimate the probability of secretory proteins with strong polyampholytes to phase separate. PSAP uses a total of 55 sequence features and employs a Random Forest method to predict the likelihood of proteins that could participate in protein phase separation **[Supplementary Table 9]**.

### Experimentally determined secretory proteins

All the experimentally determined fungal pathogenesis-related genes of plant pathogens were obtained from PHI-base (version 4.15) [59]. For these genes, we retrieved the corresponding protein sequences from Uniprot and classified them into effectors and non-effectors as described above. GO annotation of these proteins and estimation of their IDR attributes were done as described earlier **[Supplementary Table 10]**.

### Orthologous group

We retrieved information on the orthologous groups of secretory proteins from the OMA orthology database [38], referred to as OMA groups, which encompass both pairwise orthologs and hierarchical orthologous relationships [95]. Each ortho group was then classified based on its overlapping fungal class. For example, an ortho group containing orthologous proteins from nonpathogen, animal pathogen, and plant pathogen was designated as NP-AP-PP. In total, there are four distinct classes of ortho groups: NP-AP-PP, NP-AP, NP-PP, and AP-PP. We compared the IDR attributes of each protein in the ortho groups across different fungal classes in each category of orthogroups **[Supplementary Table 8]**.

### Host RNA sequencing data analysis

We curated all the RNA sequencing datasets from NCBI GEO [96] that cataloged the animal and plant host transcriptomic profiles upon different fungal infections. This search led to the identification of 30 paired-end RNA sequencing datasets spanning across 14 plant hosts and 2 animal hosts infected by 13 plant pathogens and 5 animal pathogens respectively **[Supplementary Table 11]**. FASTQ files for each dataset were retrieved and subjected to quality checks using the FastQC tool (http://www.bioinformatics.babraham.ac.uk/projects/fastqc/). Reads with a Phred score > 20 were retained for further analysis. Pre-processing, including adapter removal and filtering of low-quality reads, was performed using Cutadapt (https://doi.org/10.14806/ej.17.1.200) and Trim Galore (http://www.bioinformatics.babraham.ac.uk/projects/trim_galore/), followed by another quality check with FastQC. Sequence alignment was conducted using the Hisat2 tool [97] against the respective host reference genomes obtained from Ensembl Plants (http://plants.ensembl.org/) and Ensembl (https://www.ensembl.org/). SAMtools (version 1.11) was used for sorting, indexing, and converting SAM to BAM files. HTSeq (https://github.com/simon-anders/htseq) was employed for read counting. The expression level of genes was checked by using Fragments Per Kilobase of transcript per Million mapped reads (FPKM). Differentially expressed genes (DEGs) were identified using the DEseq2 tool from the R package, using FDR value < 0.05, calculated using the Benjamini-Hochberg correction method. Using fold change (FC) cut-offs we classified the DEGs into those that are upregulated (log_2_FC ≥ 1.5) and downregulated (log_2_FC ≤ –1.5). Each DEG was mapped to one unique protein identifier and the protein sequences of the DEGs (referred here as differentially expressed proteins; DEPs) were downloaded from the UniProt database.

### Pairwise alignments and estimation of percentage sequence identity

All strong polyampholyte IDR sequences from pathogen proteins were aligned to each strong polyampholyte IDR sequence from their respective host proteins using pairwise sequence alignment. Local alignment was performed by an in-house written R script using BLOSUM62 [98] from Biostrings package with the default gap opening penalty of 1.53 and a gap extension penalty of 0.123. Alignments with fewer than 10 aligned residues in either pathogen or host were excluded to ensure that only statistically reliable and biologically meaningful matches were considered. We next computed the sequence identity (%) between every pair of IDRs, consisting of each strong polyampholyte IDR of pathogens and each strong polyampholyte IDR of the corresponding host. The pairs with the highest sequence identity were considered for further analyses.

### Statistical analysis

We assessed the statistical significance of differences in the distributions of discrete variables using the Chi-squared test or Fisher’s exact test, while for continuous variables, we employed the non-parametric Wilcoxon rank sum test. To correct for multiple hypothesis testing, we applied the Benjamini–Hochberg method to estimate the false discovery rate (FDR). We have derived correlations between two variables using the non-parametric Spearman’s rank correlation. All statistical analyses were done using R.

## Acknowledgements

We thank P.L. Chavali and A. Dhayalan for their feedback on the manuscript. This study was supported by IISER Tirupati core funding (AM, RVK, AA, VS and SC); Ramalingaswami Re-entry Fellowship BT/RLF/Re-entry/05/2018 from Department of Biotechnology, Government of India (RVK, APS, and SC); Core Research Grant CRG/2023/004691 from Anusandhan National Research Foundation, Government of India (SC); Kishore Vaigyanik Protsahan Yojana fellowship (HR) and INSPIRE Scholarship for Higher Education (APS) from Department of Science and Technology, Government of India.

## Author contributions

The study was conceived and supervised by SC. SC and AM designed the methodology of the study. AM, RVK, AS, APS, HR, AA, and VS were involved in undertaking the analyses. AM, APS, VS and SC generated the illustrations. SC and AM wrote the original draft of the manuscript. All authors read and approved the manuscript.

## Data availability

The datasets generated and the codes written to support the findings of this study are available in the Zenodo repository (https://doi.org/10.5281/zenodo.17244321).

