## Supplementary Information for "Molecular attributes of intrinsically disordered regions in secretomes influence fungal pathogenesis"

**for**

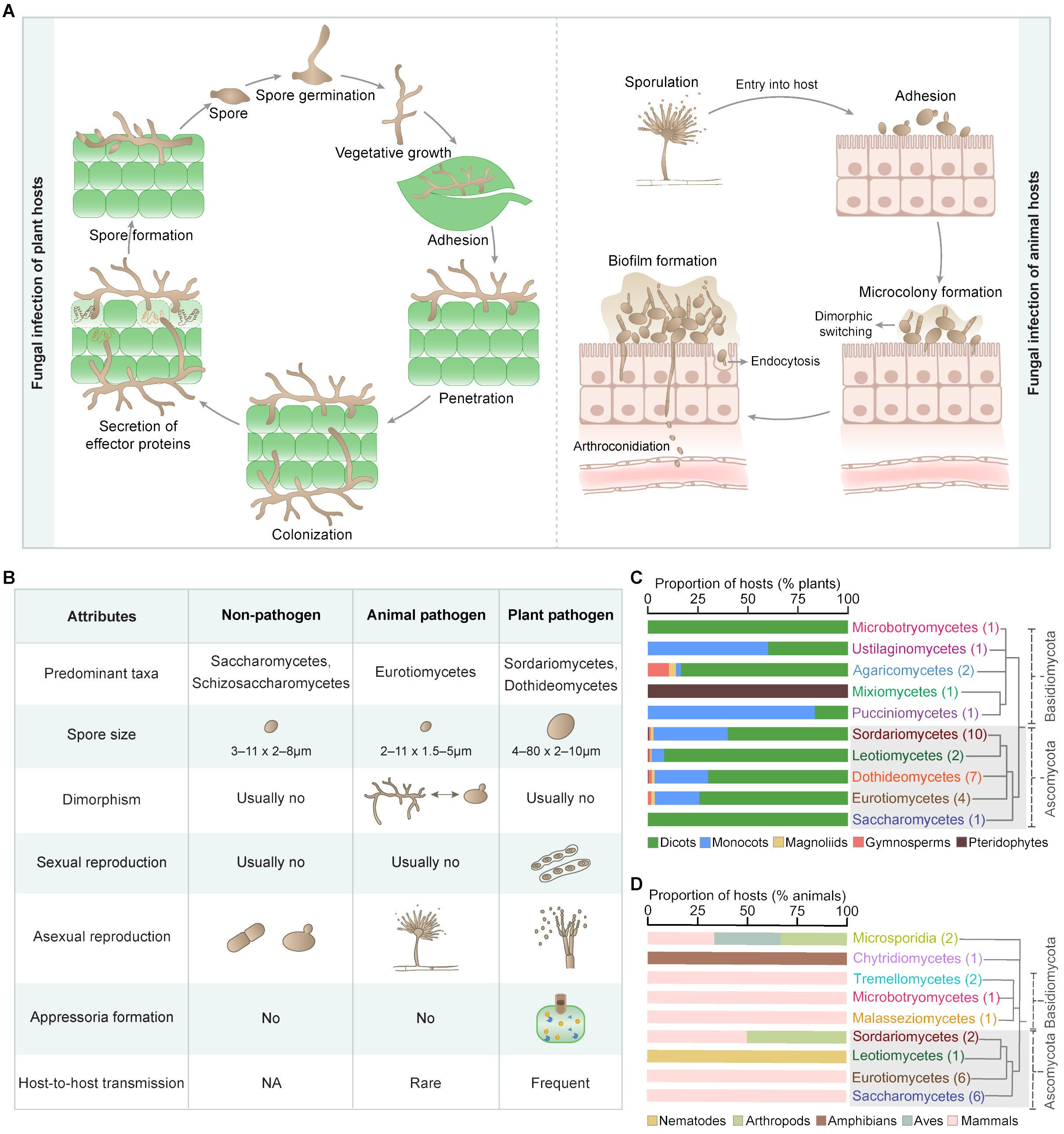


**Supplementary Fig. 1. (A)** Generalized scheme representing the different stages of plant and animal fungal infections [1-4]. **(B)** Distinct attributes of the three fungal classes studied here. Sordariomycetes and Dothideomycetes, enriched for plant pathogens, tend to produce (i) mildews and form hyphae and appressoria, (ii) sexual reproductive structures and (iii) larger spores which bring about host-to-host transmission. The ability of Eurotiomycetes, to extract nutrients by degrading the animal epidermal protein keratin explains the preponderance of animal pathogens in this taxon [1-3]. Furthermore, animal pathogens also exhibit dimorphism, whereby they switch between different morphotypes which aids in penetration of the tissues and in dissemination from a primary site of infection to other body sites [4]. Production of smaller number of spores and lack of enzymes that specifically degrade animal or plant host biomolecules may restrict the pathogenicity of Saccharomycetes and Schizosaccharomycete. Classification of **(C)** plant pathogens and **(D)** animal pathogens based on host spectrum. Bar plots depict the fraction of pathogens infecting each class of host, represented by different colors.

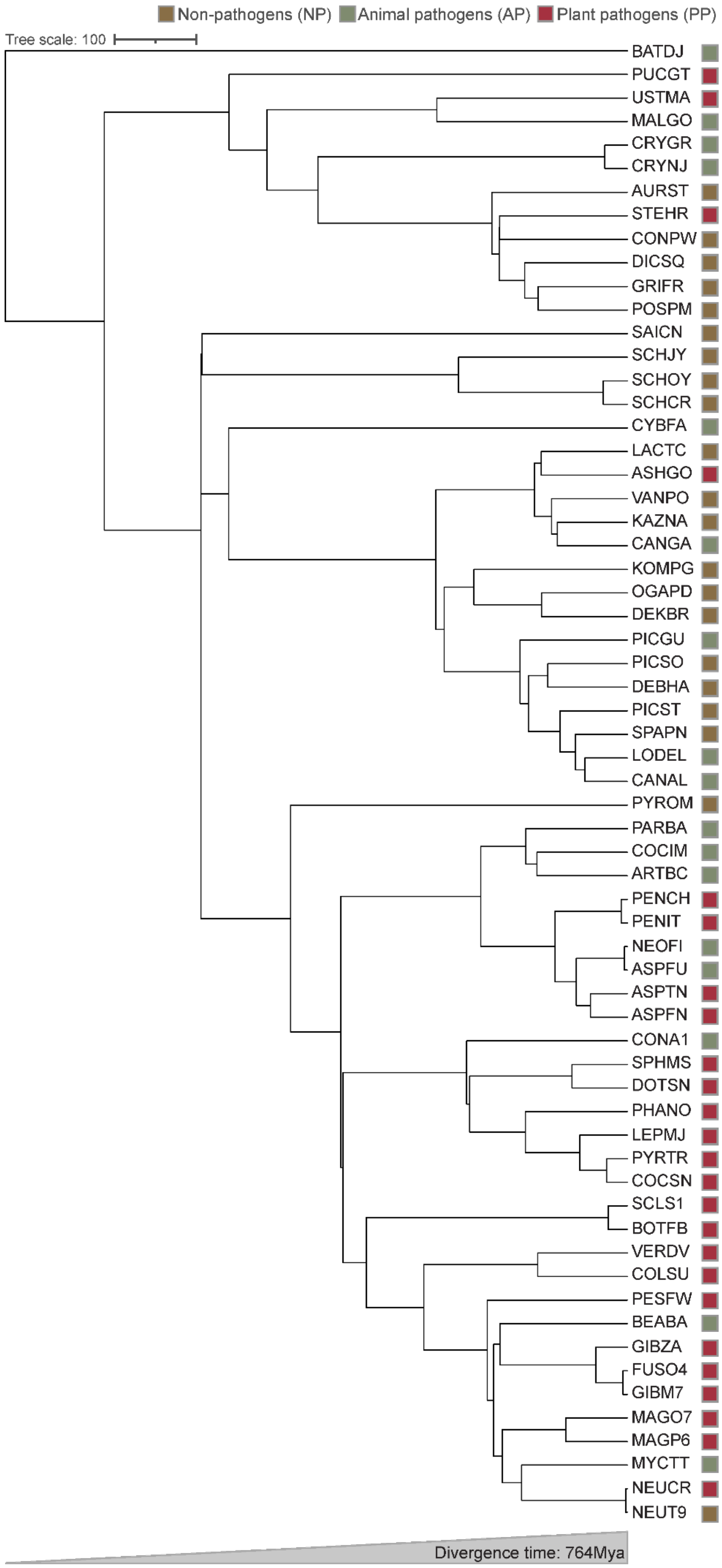
Non-pathogens (NP)
Animal pathogens (AP)
Plant pathogens (PP)
Tree scale: 100
BATDJ
PUCGT
USTMA
MALGO
CRYGR
CRYNJ
AURST
STEHR
CONPW
DICSQ
GRIFR
POSPM
SAICN
SCHJY
SCHOY
SCHCR
CYBFA
LACTC
ASHGO
VANPO
KAZNA
CANGA
KOMPG
OGAPD
DEKBR
PICGU
PICSO
DEBHA
PICST
SPAPN
LODEL
CANAL
PYROM
PARBA
COCIM
ARTBC
PENCH
PENIT
NEOFI
ASPFU
ASPTN
ASPFN
CONA1
SPHMS
DOTSN
PHANO
LEPMJ
PYRTR
COCSN
SCLS1
BOTFB
VERDV
COLSU
PESFW
BEABA
GIBZA
FUSO4
GIBM7
MAGO7
MAGP6
MYCTT
NEUCR
NEUT9
Divergence time: 764Mya

**Supplementary Fig. 2. Fungal species included in this study span a broad evolutionary timescale.** Phylogenetic tree depicts the divergence time of 63 fungal species obtained from the TimeTree database [5]. The names of the species have been abbreviated and the taxonomic and phylogenetic details of the species have been provided in **Supplementary Table 2**. The boxes adjacent to the species names depict the pathogenicity status (non-pathogen, animal or plant pathogen).

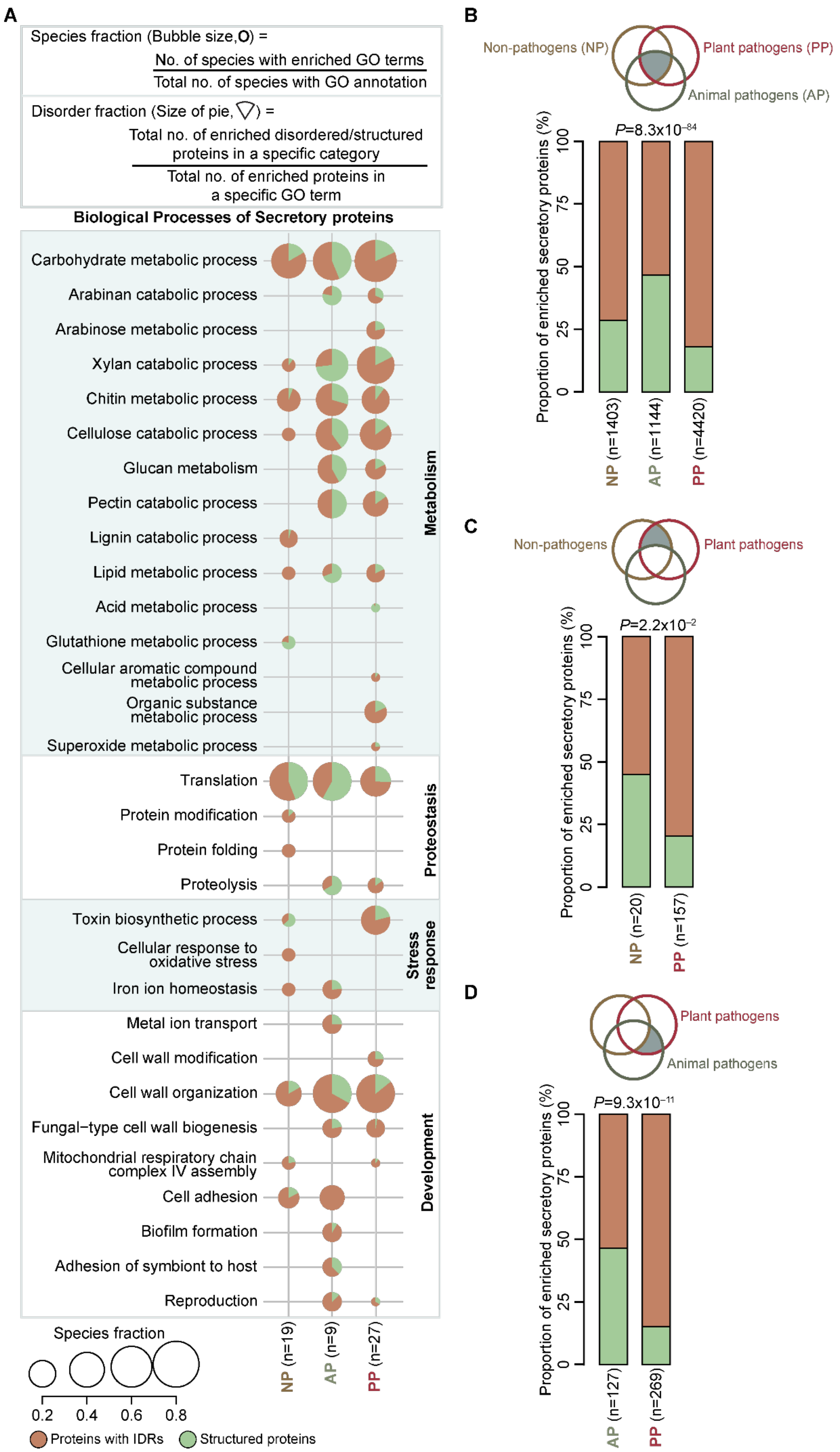
A
Species fraction (Bubble size, O) = No. of species with enriched GO terms / Total no. of species with GO annotation
Disorder fraction (Size of pie) = Total no. of enriched disordered/structured proteins in a specific category / Total no. of enriched proteins in a specific GO term
Biological Processes of Secretory proteins
Carbohydrate metabolic process
Arabinan catabolic process
Arabinose metabolic process
Xylan catabolic process
Chitin metabolic process
Cellulose catabolic process
Glucan metabolism
Pectin catabolic process
Lignin catabolic process
Lipid metabolic process
Acid metabolic process
Glutathione metabolic process
Cellular aromatic compound metabolic process
Organic substance metabolic process
Superoxide metabolic process
Metabolism
Translation
Protein modification
Protein folding
Proteolysis
Proteostasis
Toxin biosynthetic process
Cellular response to oxidative stress
Iron ion homeostasis
Stress response
Metal ion transport
Cell wall modification
Cell wall organization
Fungal−type cell wall biogenesis
Mitochondrial respiratory chain complex IV assembly
Cell adhesion
Biofilm formation
Adhesion of symbiont to host
Reproduction
Development
NP (n=19)
AP (n=9)
PP (n=27)
Species fraction
0.2
0.4
0.6
0.8
Proteins with IDRs
Structured proteins
B
Non-pathogens (NP)
Plant pathogens (PP)
Animal pathogens (AP)
P=8.3x10−84
Proportion of enriched secretory proteins (%)
0
25
50
75
100
NP (n=1403)
AP (n=1144)
PP (n=4420)
C
Non-pathogens
Plant pathogens
P=2.2x10−2
Proportion of enriched secretory proteins (%)
0
25
50
75
100
NP (n=20)
PP (n=157)
D
Plant pathogens
Animal pathogens
P=9.3x10−11
Proportion of enriched secretory proteins (%)
0
25
50
75
100
AP (n=127)
PP (n=269)

**Supplementary Fig. 3. Plant pathogens show higher fraction of IDRs in secretory proteins associated with pathogenesis-related processes. (A)** Bubble plot shows the enriched GO-Biological Processes (FDR < 0.05) manually grouped into different broad terms. The bubble size represents the number of species showing enrichment for a specific process. The pie chart depicts the proportion of structured and disordered secretory proteins. Species fraction and protein disordered/structured fraction for each enriched GO term were estimated as provided in the top panel. 'n' represents the number of species in each fungal class. Bar plot represents the distribution of the disordered and structured secretory proteins among overlapping biological processes across **(B)** all three classes of fungi, **(C)** non-pathogens and plant pathogens, as well as **(D)** animal pathogens and plant pathogens. P-value was computed using Fisher's exact test. 'n' represents the number of proteins in enriched GO terms.

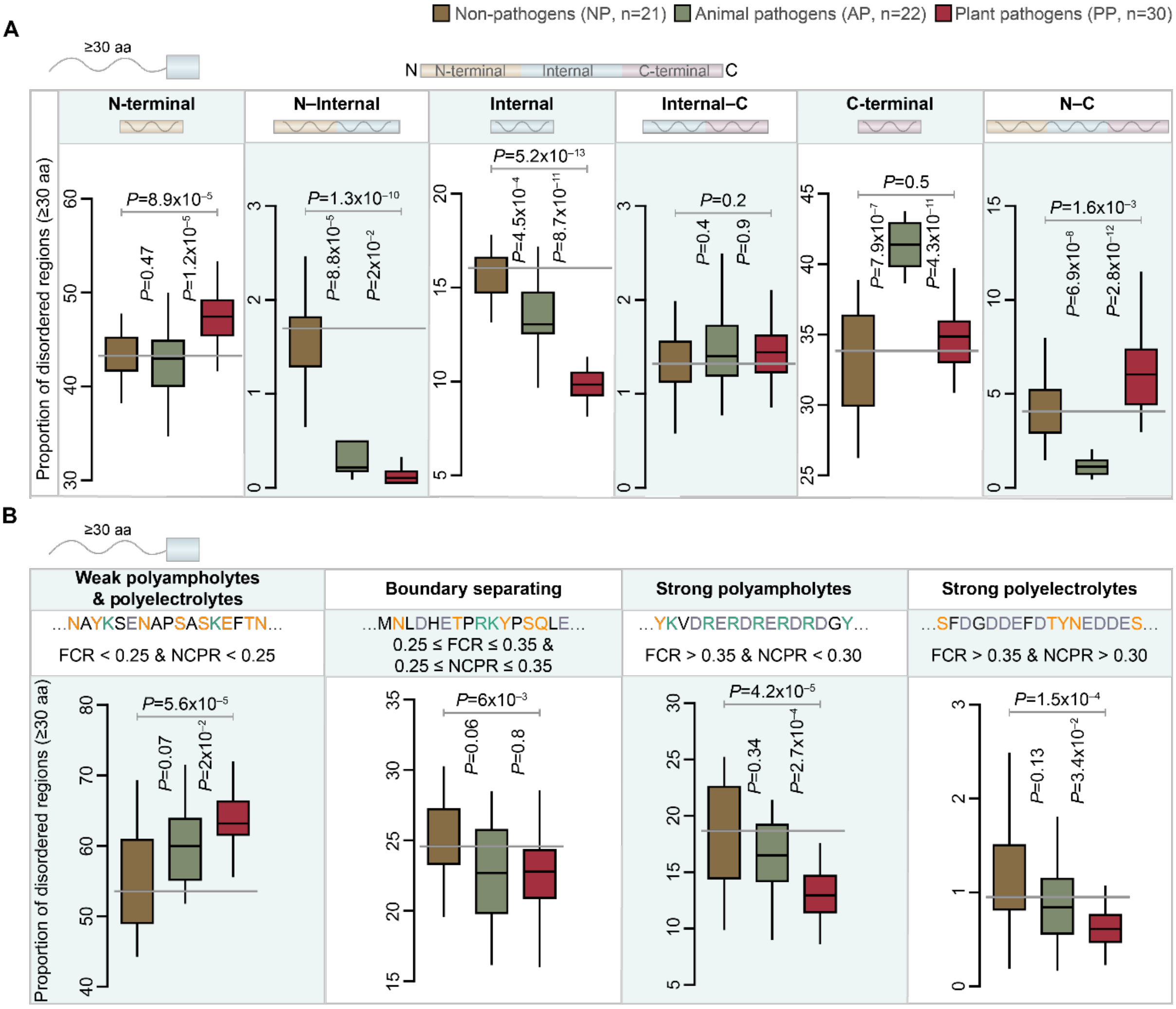


**Supplementary Fig. 4. Bias for N-terminal disordered regions and weak polyampholytes and polyelectrolytes secretory proteins with IDRs ≥ 30 amino acids long.** Boxplot showing the distribution of **(A)** disordered regions across different protein location bins, defined based on length tertile cut-offs and **(B)** different state IDRs of secretory proteins across the three classes of fungi. Statistical significance was estimated using Wilcoxon rank sum tests. Outlier datapoints have been removed to facilitate better visualization.

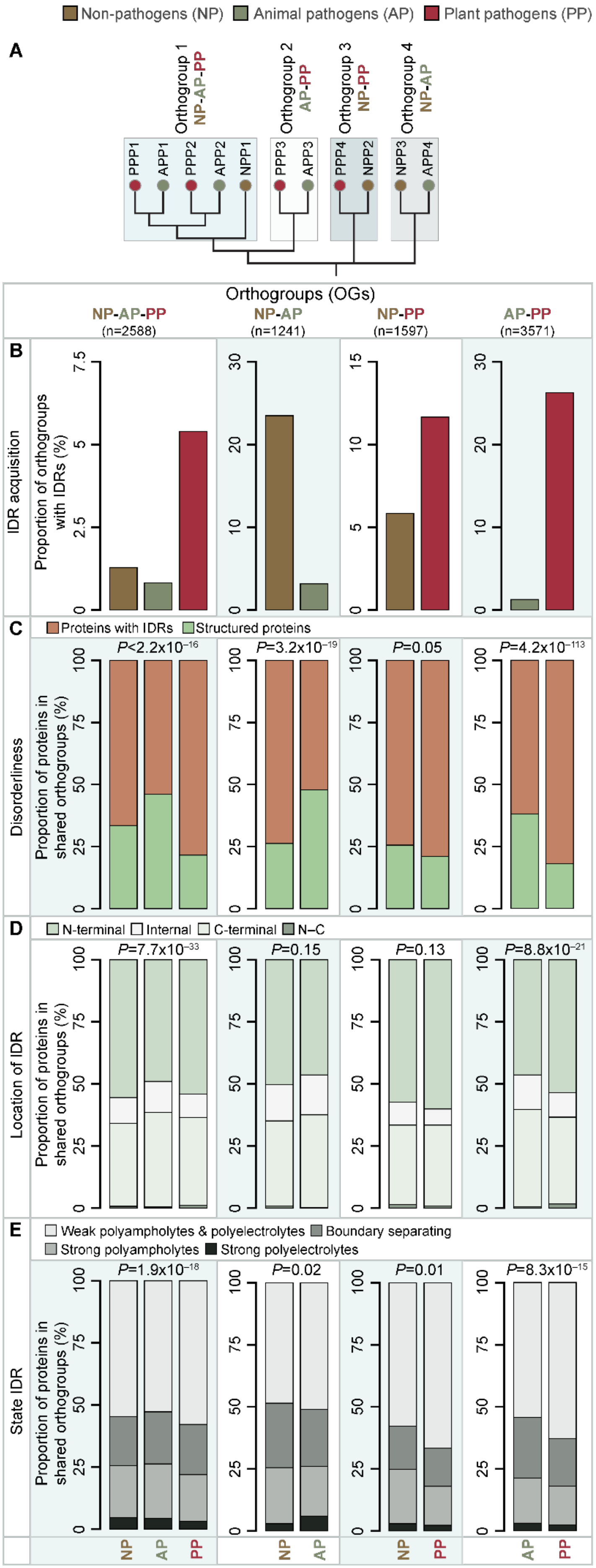
Non-pathogens (NP) Animal pathogens (AP) Plant pathogens (PP)
A
Orthogroup 1 NP-AP-PP
Orthogroup 2 AP-PP
Orthogroup 3 NP-PP
Orthogroup 4 NP-AP
PPP1 APP1 PPP2 APP2 NPP1
PPP3 APP3
PPP4 NPP2
NPP3 APP4
Orthogroups (OGs)
NP-AP-PP (n=2588)
NP-AP (n=1241)
NP-PP (n=1597)
AP-PP (n=3571)
B
IDR acquisition
Proportion of orthogroups with IDRs (%)
C
Proteins with IDRs Structured proteins
Disorderliness
Proportion of proteins in shared orthogroups (%)
P<2.2x10−16
P=3.2x10−19
P=0.05
P=4.2x10−113
D
N-terminal Internal C-terminal N–C
Location of IDR
P=7.7x10−33
P=0.15
P=0.13
P=8.8x10−21
E
Weak polyampholytes & polyelectrolytes Boundary separating
Strong polyampholytes Strong polyelectrolytes
State IDR
P=1.9x10−18
P=0.02
P=0.01
P=8.3x10−15
NP AP PP
NP AP
NP PP
AP PP

**Supplementary Fig. 5. Orthologs of plant pathogen secretory proteins show acquisition of IDR attributes that might facilitate pathogenesis. (A)** Classification of shared orthologous groups (orthogroups obtained from OMA database [6]) into three categories– those that contain proteins from all three (NP-AP-PP) or any two fungal classes (NP-AP, NP-PP and AP-PP), where PPP represents plant pathogen protein, APP represents animal pathogen protein and NPP represents non-pathogen protein. **(B)** Proportion of different categories of orthogroups that have shown acquisition of IDRs in only one fungal class (NP, AP or PP) while the rest of the orthologs are structured. In the NP-AP-PP, NP-PP and AP-PP categories, plant pathogen orthogroups show higher proportion of acquisition of IDRs compared with other classes of fungi, while among NP-AP, non-pathogens tend to show higher IDR acquisition. **(C)** Bar plot showing the proportion of proteins with IDRs in different categories of orthogroups. In orthogroups which contained proteins with IDRs across the fungal classes, we compared the proportion of proteins with IDRs of each fungal class with the other(s) in that orthogroup. Similar to our observations in panel **(B)**, we found that plant pathogen proteins tend to be more disordered. **(D)** Bar plot of distribution of the IDRs across the four protein length bins. Overlapping bins (N–Internal, Internal-C) were disregarded due to fewer data points. IDRs in the orthologous proteins of plant pathogens are preponderant in N-terminal region and that of animal pathogens show preponderance in C-terminal regions. **(E)** Bar plot displaying the proportion of different state IDRs across proteins from different fungal classes. Plant pathogen proteins show higher proportion of weak polyampholyte and polyelectrolyte IDRs.

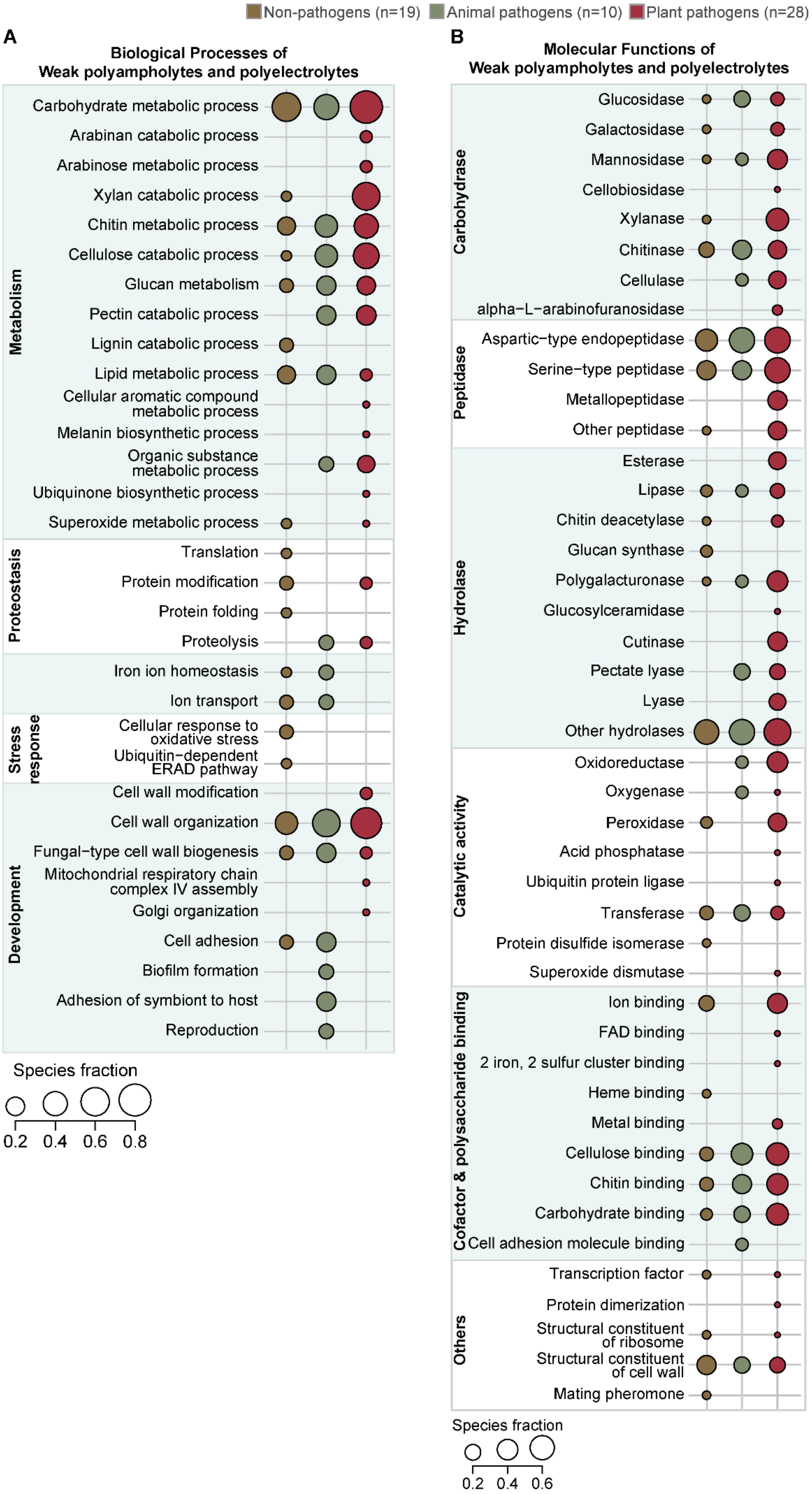
Non-pathogens (n=19)
Animal pathogens (n=10)
Plant pathogens (n=28)
A
Biological Processes of
Weak polyampholytes and polyelectrolytes
Metabolism
Carbohydrate metabolic process
Arabinan catabolic process
Arabinose metabolic process
Xylan catabolic process
Chitin metabolic process
Cellulose catabolic process
Glucan metabolism
Pectin catabolic process
Lignin catabolic process
Lipid metabolic process
Cellular aromatic compound metabolic process
Melanin biosynthetic process
Organic substance metabolic process
Ubiquinone biosynthetic process
Superoxide metabolic process
Proteostasis
Translation
Protein modification
Protein folding
Proteolysis
Iron ion homeostasis
Ion transport
Stress response
Cellular response to oxidative stress
Ubiquitin−dependent ERAD pathway
Development
Cell wall modification
Cell wall organization
Fungal−type cell wall biogenesis
Mitochondrial respiratory chain complex IV assembly
Golgi organization
Cell adhesion
Biofilm formation
Adhesion of symbiont to host
Reproduction
Species fraction
0.2 0.4 0.6 0.8
B
Molecular Functions of
Weak polyampholytes and polyelectrolytes
Carbohydrase
Glucosidase
Galactosidase
Mannosidase
Cellobiosidase
Xylanase
Chitinase
Cellulase
alpha−L−arabinofuranosidase
Peptidase
Aspartic−type endopeptidase
Serine−type peptidase
Metallopeptidase
Other peptidase
Hydrolase
Esterase
Lipase
Chitin deacetylase
Glucan synthase
Polygalacturonase
Glucosylceramidase
Cutinase
Pectate lyase
Lyase
Other hydrolases
Catalytic activity
Oxidoreductase
Oxygenase
Peroxidase
Acid phosphatase
Ubiquitin protein ligase
Transferase
Protein disulfide isomerase
Superoxide dismutase
Cofactor & polysaccharide binding
Ion binding
FAD binding
2 iron, 2 sulfur cluster binding
Heme binding
Metal binding
Cellulose binding
Chitin binding
Carbohydrate binding
Cell adhesion molecule binding
Others
Transcription factor
Protein dimerization
Structural constituent of ribosome
Structural constituent of cell wall
Mating pheromone
Species fraction
0.2 0.4 0.6

**Supplementary Fig. 6. Secretory proteins with weak polyampholytes and polyelectrolytes are enriched for carbohydrate metabolism-related processes.** Bubble plot indicating the number of significantly enriched (FDR<0.05) Gene Ontology **(A)** Biological Processes and **(B)** Molecular Functions, which have been manually grouped into broad terms. The bubble size denotes the fraction of species (estimated as shown in **Supplementary Fig. 1**) in which the respective GO terms are enriched.

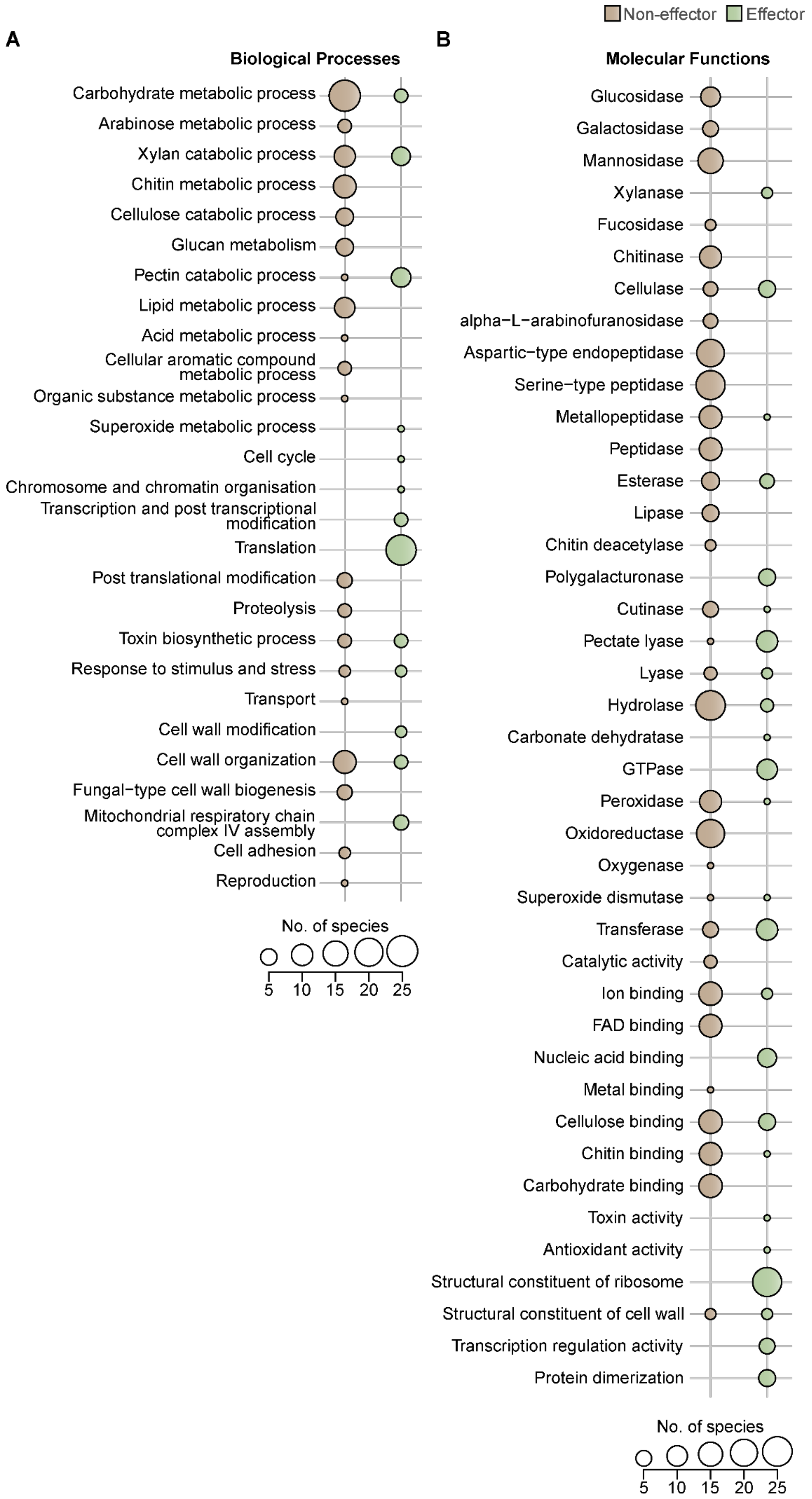
Non-effector
Effector
A
Biological Processes
Carbohydrate metabolic process
Arabinose metabolic process
Xylan catabolic process
Chitin metabolic process
Cellulose catabolic process
Glucan metabolism
Pectin catabolic process
Lipid metabolic process
Acid metabolic process
Cellular aromatic compound metabolic process
Organic substance metabolic process
Superoxide metabolic process
Cell cycle
Chromosome and chromatin organisation
Transcription and post transcriptional modification
Translation
Post translational modification
Proteolysis
Toxin biosynthetic process
Response to stimulus and stress
Transport
Cell wall modification
Cell wall organization
Fungal−type cell wall biogenesis
Mitochondrial respiratory chain complex IV assembly
Cell adhesion
Reproduction
No. of species
5 10 15 20 25
B
Molecular Functions
Glucosidase
Galactosidase
Mannosidase
Xylanase
Fucosidase
Chitinase
Cellulase
alpha−L−arabinofuranosidase
Aspartic−type endopeptidase
Serine−type peptidase
Metallopeptidase
Peptidase
Esterase
Lipase
Chitin deacetylase
Polygalacturonase
Cutinase
Pectate lyase
Lyase
Hydrolase
Carbonate dehydratase
GTPase
Peroxidase
Oxidoreductase
Oxygenase
Superoxide dismutase
Transferase
Catalytic activity
Ion binding
FAD binding
Nucleic acid binding
Metal binding
Cellulose binding
Chitin binding
Carbohydrate binding
Toxin activity
Antioxidant activity
Structural constituent of ribosome
Structural constituent of cell wall
Transcription regulation activity
Protein dimerization
No. of species
5 10 15 20 25

**Supplementary Fig. 7. Plant pathogen non-effectors and effectors are enriched for distinct pathogenesis-related processes.** Bubble plot indicating the number of significantly enriched (FDR<0.05) Gene Ontology **(A)** Biological Processes and **(B)** Molecular Functions that are manually grouped into broad terms. The bubble size denotes the number of species representing enriched GO terms in each category.

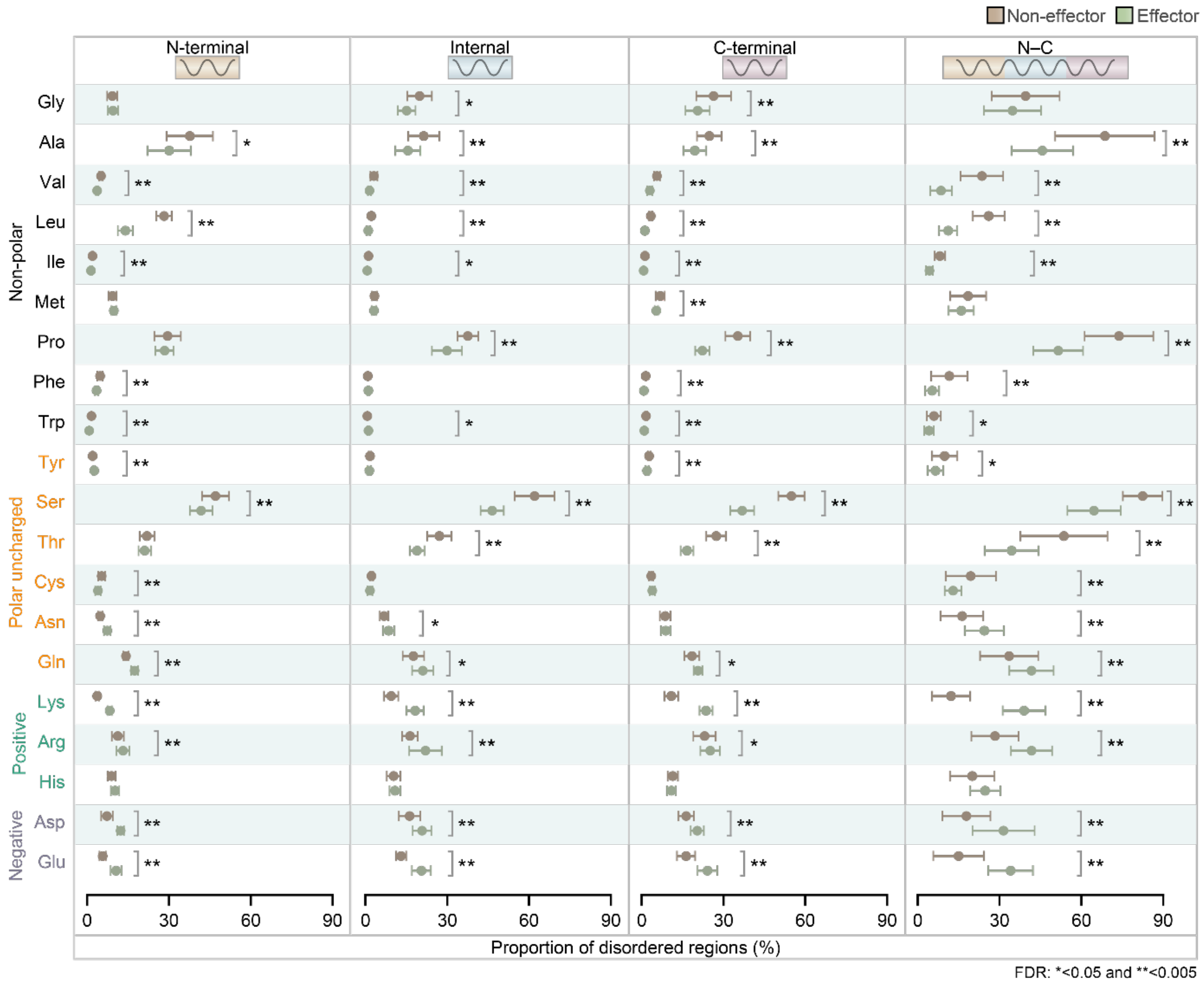


**Supplementary Fig. 8. Effectors show higher proportion of charged amino acids in IDRs.** Distribution of amino acids among IDRs of non-effectors and effectors spanning different protein location bins, which were defined using protein length tertiles. For each amino acid circle represents the median value, the bars represent Median Absolute Deviation (MAD) and the color indicates the class of plant pathogen secretory proteins. Statistical significance was estimated using Wilcoxon rank sum test, and asterisks near the bars indicate the False Discovery Rates (FDR) computed using Benjamini-Hochberg method of correction for multiple testing.

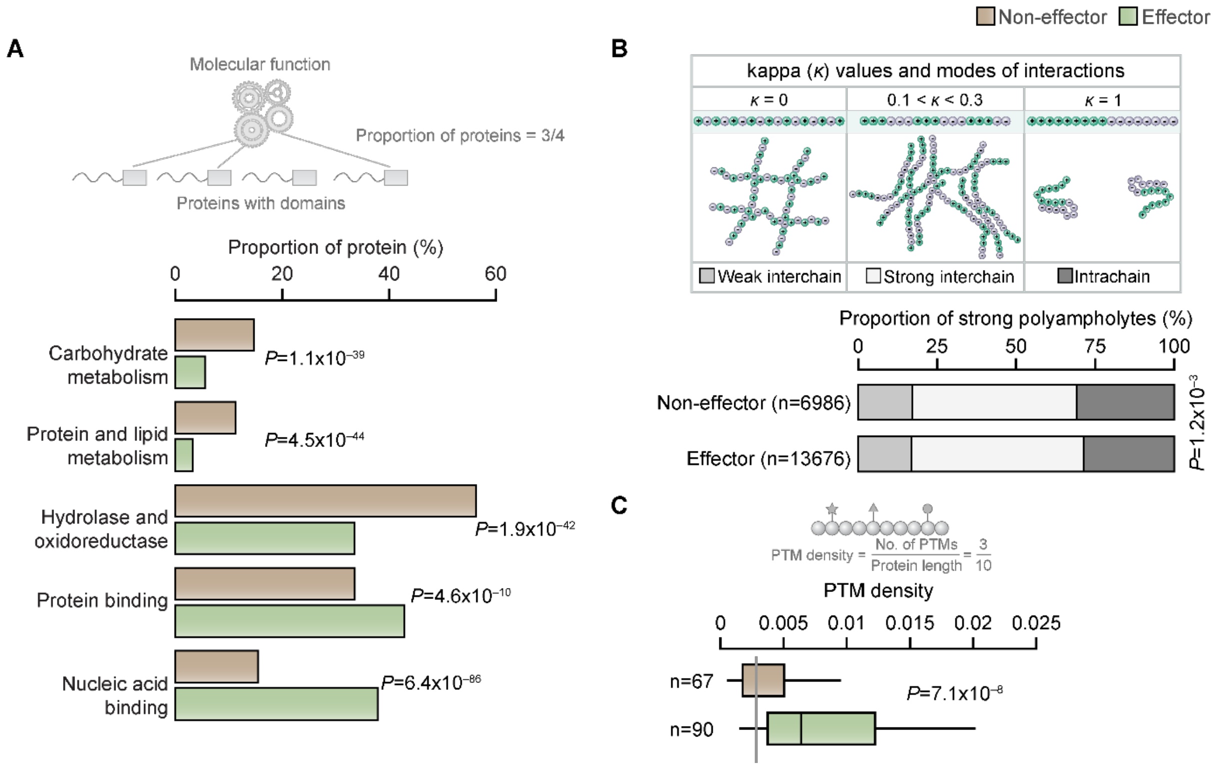


**Supplementary Fig. 9. Effectors predominantly contain functional regions involved in transcriptional and translational regulation. (A)** Bar plot representing the proportion of non-effectors and effectors with N-terminal weak polyampholyte and polyelectrolyte containing proteins across various GO Molecular Functions annotated for protein domains present in them. **(B)** Bar plot represents the fraction of proteins with strong polyampholytes that bring about different types of interactions, classified based on charge patterning as estimated by kappa values; Intrachain ($\kappa \leq 0.1$), Strong interchain ($0.1<\kappa<0.3$), Weak interchain ($\kappa \geq 0.3$). Charge patterning is a key feature that regulates the conformation, number of binding partners and function of strong polyampholytes [7]. A κ value near zero (well mixed oppositely charged residues) facilitates limited protein-protein interactions (PPIs) through weak interchain interaction while a κ value tending towards 1 (completely segregated charges) promotes intrachain interactions and does not support PPIs. However, κ values ranging from 0.1 to 0.3 promotes PPIs and a higher number of interchain interactions. A higher number of interchain interactions tend to facilitate formation of phase-separated condensates [8]. Statistical significance was estimated using Fisher's exact test. **(C)** Box plot depicting the distribution of post-translational modification (PTM) density (no. of PTMs normalized by the protein length) among effectors and non-effectors containing strong polyampholytes. Statistical significance was computed using Wilcoxon rank sum tests. 'n' denotes the number of proteins in each class.

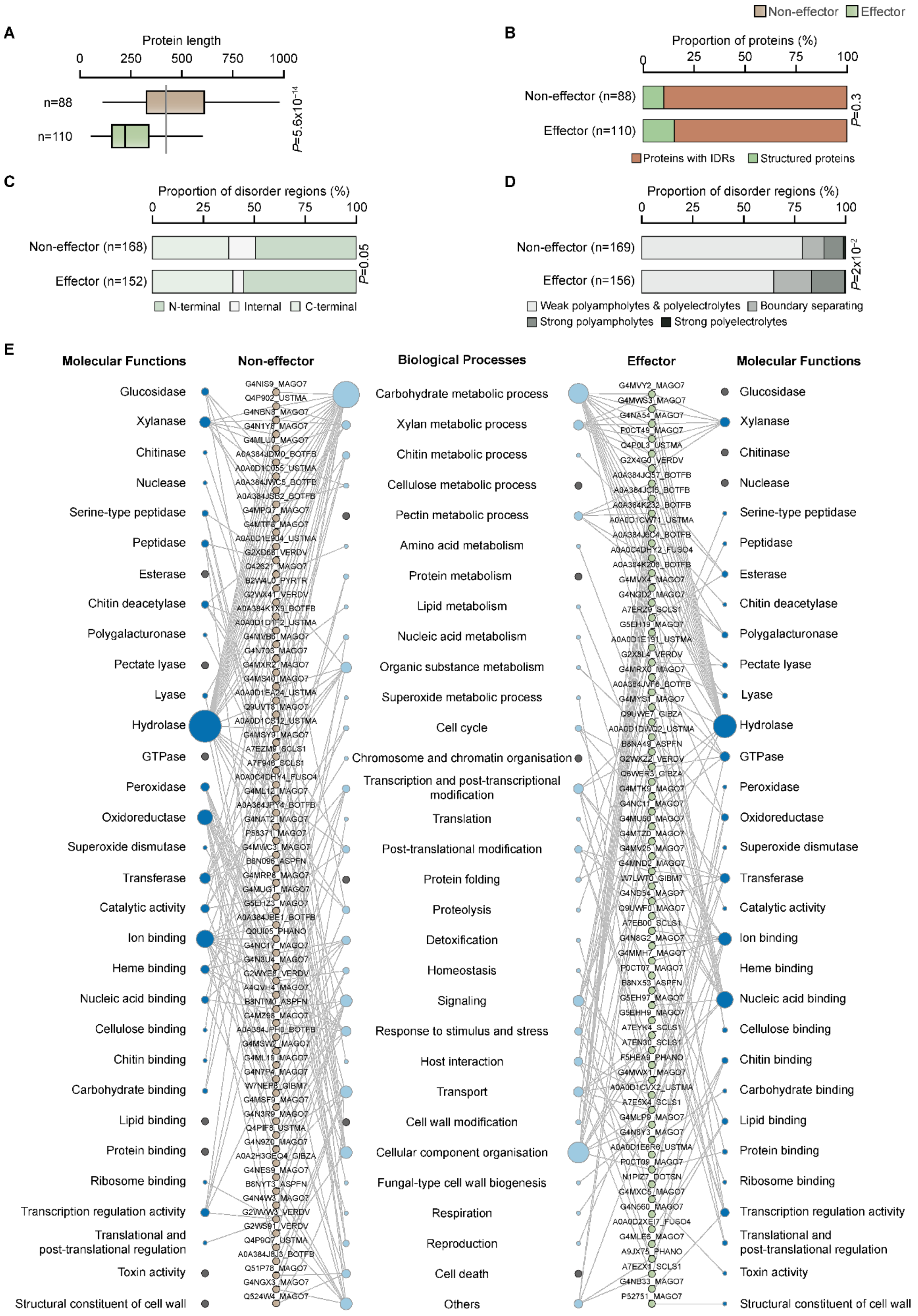
A
Protein length
0 250 500 750 1000
n=88
n=110
P=5.6x10^-14
B
Non-effector
Effector
Proportion of proteins (%)
0 25 50 75 100
Non-effector (n=88)
Effector (n=110)
P=0.3
Proteins with IDRs
Structured proteins
C
Proportion of disorder regions (%)
0 25 50 75 100
Non-effector (n=168)
Effector (n=152)
P=0.05
N-terminal
Internal
C-terminal
D
Proportion of disorder regions (%)
0 25 50 75 100
Non-effector (n=169)
Effector (n=156)
P=2x10^-2
Weak polyampholytes & polyelectrolytes
Boundary separating
Strong polyampholytes
Strong polyelectrolytes
E
Molecular Functions
Non-effector
Biological Processes
Effector
Molecular Functions
Glucosidase
Xylanase
Chitinase
Nuclease
Serine-type peptidase
Peptidase
Esterase
Chitin deacetylase
Polygalacturonase
Pectate lyase
Lyase
Hydrolase
GTPase
Peroxidase
Oxidoreductase
Superoxide dismutase
Transferase
Catalytic activity
Ion binding
Heme binding
Nucleic acid binding
Cellulose binding
Chitin binding
Carbohydrate binding
Lipid binding
Protein binding
Ribosome binding
Transcription regulation activity
Translational and post-translational regulation
Toxin activity
Structural constituent of cell wall
Carbohydrate metabolic process
Xylan metabolic process
Chitin metabolic process
Cellulose metabolic process
Pectin metabolic process
Amino acid metabolism
Protein metabolism
Lipid metabolism
Nucleic acid metabolism
Organic substance metabolism
Superoxide metabolic process
Cell cycle
Chromosome and chromatin organisation
Transcription and post-transcriptional modification
Translation
Post-translational modification
Protein folding
Proteolysis
Detoxification
Homeostasis
Signaling
Response to stimulus and stress
Host interaction
Transport
Cell wall modification
Cellular component organisation
Fungal-type cell wall biogenesis
Respiration
Reproduction
Cell death
Others

**Supplementary Fig. 10. Effectors and non-effector among experimentally identified pathogenesis-related proteins show trends similar to those observed for predicted effectors and non-effectors.** **(A)** Box plot of distribution of the length of experimentally identified effectors and non-effectors. Statistical significance was estimated using Wilcox rank sum test. Bar plot of distribution of **(B)** proteins with IDRs and structured proteins, **(C)** IDRs spanning different location bins and **(D)** different state IDRs of effectors and non-effectors. Statistical significance was computed using Fisher's exact test. n in panels **(A)** and **(B)** represent the number of proteins, while that in panels **(C)** and **(D)** indicate the number of IDRs. In panel **(C)** only predominant location bins (with atleast 5 instances) have been represented. **(E)** Network representation of the GO Biological Processes and Molecular Functions of experimentally determined effectors and non-effectors, recapitulates the enriched GO terms observed for predicted effectors and non-effectors. GO annotations were manually categorized into broad classes. Since the number of experimentally identified proteins were less, we could not perform GO term enrichment analysis. The bubble size denotes the number of annotated proteins. GO terms which lack representations in effectors or non-effectors are highlighted using grey nodes.

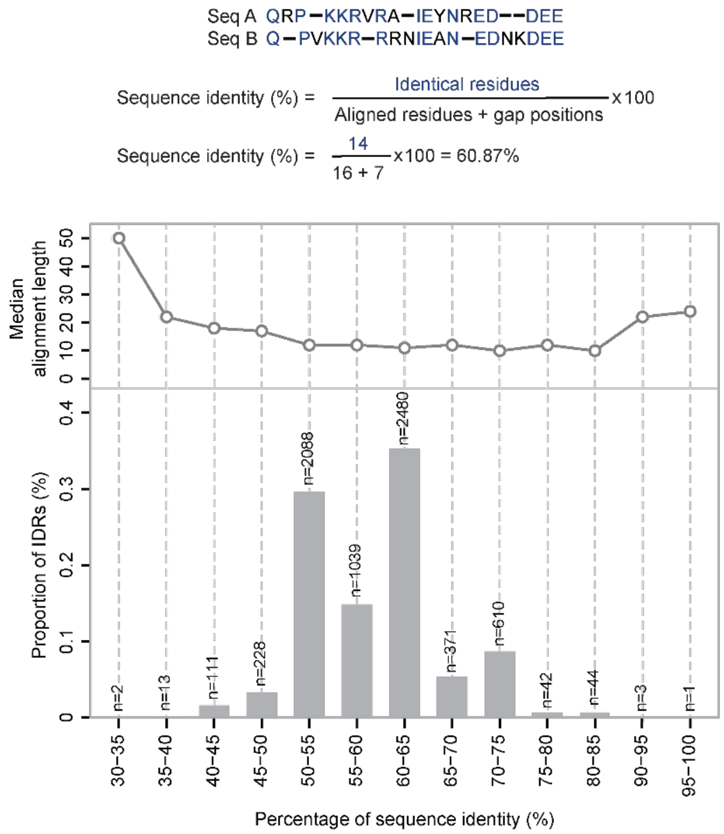


**Supplementary Fig. 11. Pairwise sequence alignment of strongly polyampholyte containing IDRs from plant pathogens and their corresponding host proteins suggests possible convergent evolution.** Percentage sequence identity was estimated by considering both aligned positions and gaps in the pairwise alignment of each plant pathogen IDR with strong polyampholyte with the corresponding host DEP IDR with strong polyampholyte (top panel). The bar plot at the bottom shows the highest percentage sequence identity for each pathogen IDR and host DEP IDR with strong polyampholytes. For this, we first generated pairwise sequence alignment for each plant pathogen IDR with strong polyampholyte against all the corresponding host DEP IDRs with strong polyampholyte and then selected the pairwise sequence alignment with the highest percentage sequence identity. For pathogens with more than one host, we selected the pairwise sequence alignment with the highest percentage sequence identity across all the host DEP IDRs. ‘n’ represents the number of pathogen IDRs with strong polyampholytes that correspond to each percentage sequence identity bin. The line plot represents the median alignment length for sequence pairs within each percentage sequence identity bin.
